# When label-free morphology is sufficient for targeted cellular measurements

**DOI:** 10.64898/2026.09.22.753654

**Authors:** Mengran Li, Jianqing Zhu, Bo Li, Chengyang Zhang, Zhenchao Tang, Jiaying Wang, Wenbin Xing, Boyu Zhang, Jinfeng Xu, Lingbei Meng, Bob Zhang, Junzhou Chen, Ronghui Zhang, Lian Zhang, Jinchao Xu

## Abstract

How cells respond to genetic perturbations reveals gene function, but each imaging experiment can measure only a limited set of molecular markers. Predicting additional readouts from label-free images could expand what these experiments reveal, but when such predictions can replace direct measurements remains unclear. Here we introduce MorphoSuff, a framework for assessing whether predicted cellular readouts support specific biological tasks. It evaluates reference reproducibility and repeats the intended analyses with predicted rather than measured responses. Rules applied to calibration perturbations select uses for testing on separate, held-out perturbations. Using nearly 10 million same-cell phase–reporter observations spanning 1,000 gene knock-outs, 52 fluorescent reporters and 73 A549 screens, we find information about reporter states beyond average perturbation responses and cell size, shape and density. Disrupting same-cell pairing within perturbation and screen groups during training reduced the median correlation between predicted and measured cell phenotypes from 0.76 to 0.29. Predictions often preserved perturbation rankings while compressing response magnitudes. Within the atlas, ranking-based acceptance supported more successful prioritization uses than acceptance based on Pearson correlation alone. In primary hepatocytes, brightfield predictions recovered 10 of the 11 strongest metabolic-activity losses among 217 held-out compounds, compared with 8 of 11 for a matched Ridge predictor using fluorescence-derived cell counts. MorphoSuff evaluates predictors by the scientific uses they support, using reference reproducibility and held-out task outcomes to assess whether inferred measurements are adequate.

## Introduction

Genetic screens reveal gene function by recording how perturbations change the state of a cell ^1^, but optical channels, reagents and throughput limit each experiment to part of that state. Imaging screens capture organelle organization, morphology and selected molecular markers ^2,3^. Perturb-seq measures changes in gene expression ^4,5^, and Perturb-CITE-seq adds cell-surface proteins ^6^. AI-powered virtual-cell models aim to connect these different observations with predictions of cellular responses ^7^. Used as virtual instruments, they could supply readouts that an experiment did not collect. Such predictions are useful when they preserve the information needed for the intended biological analysis. When can a predicted readout stand in for a direct measurement?

Label-free cell imaging offers a practical setting in which to test this possibility. Phase images capture cellular appearance without target-specific labels and can be collected repeatedly ^8^. Pooled optical screening connects each imaged cell with its perturbation ^9^. Recent platforms extend image-based genetic screening across large perturbation collections ^10,11^ and to primary cells and tissues ^12^. Quantitative image analysis converts cellular appearance into profiles of morphology and subcellular organization ^13^. These profiles reveal similarities among perturbations ^14^ and predict measures of cell health ^15^. If label-free images could also stand in for targeted readouts, the same acquisition could support several biological analyses without additional target-specific labels.

Previous work has inferred fluorescent labels from unlabelled images ^16,17^, including channel prediction for high-content screening ^18^. Related approaches connect images with molecular profiles ^19,20^ or infer surface proteins from RNA ^21^. Morphological and transcriptional perturbation profiles contain both shared and complementary information ^22,23^. These capabilities concern different properties of a cellular readout: whether it distinguishes meaningful perturbations, shares response patterns with morphology, or can be predicted for an unseen perturbation. Whether a prediction can replace that readout in a given analysis is a further question. The requirements depend on the task. A follow-up screen may need only to prioritize a few strong perturbations; comparing effect sizes additionally requires accurate response magnitudes, and identifying affected pathways requires preservation of the relevant functional analysis.

Three challenges complicate the use of predicted readouts as measurements. First, agreement can be driven by average responses of perturbation groups or simple properties such as cell count ^24^; it does not by itself establish recovery of individual-cell states. Second, a learned relationship can change with cellular context ^25^, so performance in one screen may not persist in another. Third, the measured reference must itself be reproducible enough to judge a substitute. Application-appropriate validation ^26^ therefore requires assessing reference reproducibility, the response properties retained by prediction and the outcome of the intended biological analysis.

Here we introduce MorphoSuff, a framework for assessing *measurement sufficiency*: whether replacing a measured readout with a prediction preserves the evidence required for a specified biological task. Each assessment concerns a particular readout, predictor, experimental context and intended use. Reference reproducibility and response fidelity characterize the inferred measurement; repeating the intended analysis with predicted rather than measured responses tests whether it supports perturbation prioritization, effect-size comparison or functional interpretation. A paired pilot provides data for fitting a predictor and assessing its possible uses. Fixed acceptance rules applied to calibration perturbations select uses for testing on separate evaluation perturbations, where successes, failures and missed opportunities are counted. We evaluate this approach in an optical CRISPR atlas of 1,000 gene knockouts, 52 fluorescent reporters and 73 screens in A549 lung cancer cells ^27^. Nearly 10 million same-cell phase–reporter observations span ten biological systems, including subcellular organization, signalling and cellular stress. This breadth supports comparisons of individual-cell information, response properties and retained functional evidence across diverse target states under a common evaluation design. Independent hepatocyte, cell-health and RNA datasets test specific uses across different readouts and pairing designs; Cell Health and LINCS additionally test whether calibration evidence selects uses that succeed on held-out perturbations. By evaluating both the adequacy of predicted measurements for a task and the reliability of selecting that use, MorphoSuff connects model validation to the scientific analyses that predictions can support.

## Results

MorphoSuff evaluates both what an inferred readout preserves and whether that evidence supports its use in a specified analysis. In the atlas, we tested prioritization of strong perturbation responses on evaluation genes held apart from calibration genes, using fixed MLP predictions. Across 260 reporter–fold cases, acceptance based on response-magnitude ranking supported 204 successful uses with seven failures, compared with 184 successes and eight failures for acceptance based on Pearson recovery alone (Fig. 3e,f). Combining response criteria selected a smaller set of 170 successful uses with one failure. These comparisons quantify how validation criteria change the uses supported and the failures encountered.

### A paired atlas links morphology to targeted cellular phenotypes

Pooled optical screening links each cell’s phase morphology, reporter phenotype and sgRNA identity in a common record (Fig. 1a). These paired observations allow us to assess prediction of cellular phenotypes and reconstruction of perturbation responses. We then use reference reproducibility and response fidelity to characterize the inferred readout, and evaluate whether it supports a specified biological task on held-out perturbations.

**Figure 1.**
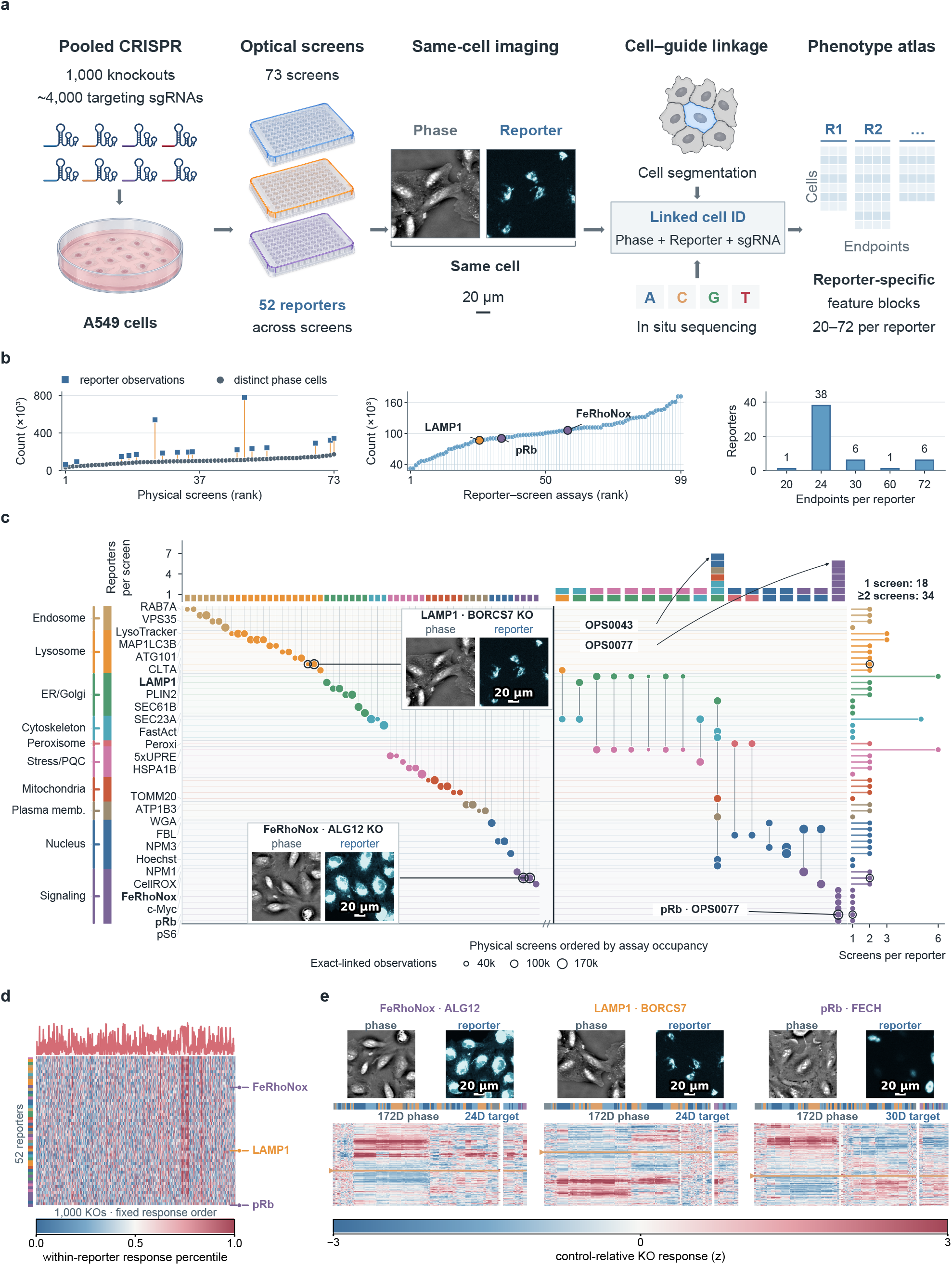
Measurement sufficiency assessed in a paired perturbation atlas. **a**, Pooled CRISPR screens and linkage of phase, reporter and sgRNA measurements to the same cell. Drawings are schematic; microscopy shows an exact-linked phase–reporter pair. **b**, Counts across 73 screens, coverage of 99 assays and reporter frequencies by endpoint-block size: 7,344,374 phase cells, 9,996,286 phase–reporter observations and 1,604 endpoints. **c**, Reporter–screen topology; blanks denote unacquired assays and marginal tracks show recurrence, occupancy and depth. **d**, Control-relative knockout responses, converted to within-reporter percentiles and ordered by recoverability. **e**, Exact-linked phase/reporter crops and response maps for FeRhoNox, LAMP1 and pRb. Images use reporter-specific contrast; scale bars, 20 *µ*m.

Quality-controlled atlas linkage yielded 7,344,374 phase cells and 9,996,286 phase–reporter observations: one cell could carry multiple reporter measurements. Each cell had 172 engineered phase features, paired with 20–72 fluorescence-derived endpoints describing intensity and spatial organization. The 52 reporters contributed 1,604 endpoints across ten biological systems (Fig. 1b).

The acquisition graph connected these reporters to 73 screens through 99 acquired assays; unacquired combinations were missing assays, not negative observations (Fig. 1c). Each screen contained one to seven reporters, and 34 reporters supported destination-screen testing. Unlike collections repeating a common panel ^10,28^, this topology distinguishes reporter, screen and assay summaries: a screen average reflects both its reporter composition and its acquisition context. The reporters also differ in the phenotypes they measure, so the 1,604 endpoints form reporter-specific blocks rather than one common measurement panel. Reporter programmes and exact-linked FeRhoNox, LAMP1 and pRb examples show this diversity of reference responses (Fig. 1d,e).

### Predictor choice changes recovery across generalization settings

We compared ten methods spanning reporter-specific tabular prediction, shared multitask prediction and multimodal completion (Methods). These approaches include models fitted to one reporter at a time, models trained jointly across reporters and methods developed to predict one measurement modality from another. All received the same phase representation, target endpoints and predefined partitions.

The evaluation separated three scientific questions: prediction in an observed perturbation context, reconstruction of responses to unseen knockout genes, and transfer to a new acquisition environment. Field, gene and whole-screen holdouts tested these questions, respectively (Fig. 2a). Cell correlations measured exact-linked phenotype recovery, whereas knockout correlations measured re-construction of control-relative perturbation effects. Reporter, screen and reporter–screen summaries then located where recovery was retained. Both the evaluation target and reporting unit can affect method rankings ^29,30^.

**Figure 2.**
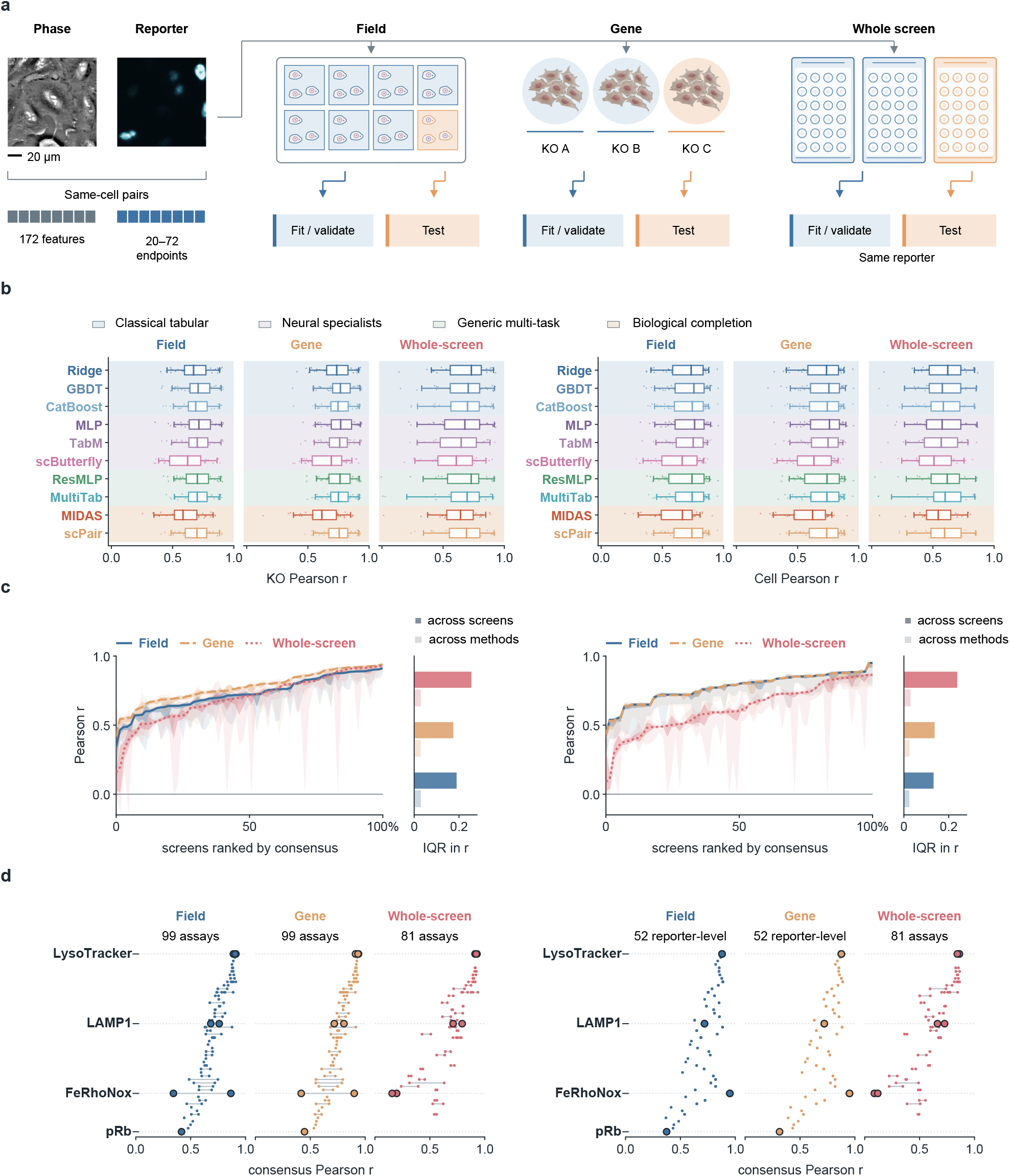
Benchmark performance across generalization, evaluation and aggregation axes. **a**, Exact-linked phase–reporter inputs and three schematic group holdouts: complete imaging fields, knockout genes and whole screens. Blue denotes fitting and validation data; orange denotes held-out data. Reporter identity is retained across screens; scale bar, 20 *µ*m. **b**, Reporter-level Pearson correlations for ten methods, averaged over five predefined partitions or directed destination-screen evaluations; b– d show knockout responses on the left and cell phenotypes on the right. Points denote reporters; boxes show medians and interquartile ranges, whiskers 5th–95th percentiles and pale bands method families. **c**, Physical screens ranked within each test by the unweighted median across methods, shown as lines with interquartile (dark) and full-range (pale) bands. Adjacent bars compare interquartile variation across screens (solid) and across methods within screens (outlined). **d**, Reporter–screen assays in a shared reporter order; each point denotes an assay and connectors span assays for the same reporter. Field/gene tests contain 52 reporters, 73 screens and 99 assays; whole-screen transfer contains 34 reporters, 66 destination screens and 81 directed assays. Whole-screen cell values are evaluated directly in destination cells. For field/gene cell-level screen and assay views, common per-cell predictions were unavailable for every method; these descriptive panels reweight frozen reporter-level macro-Pearson values by observed screen or assay membership. The 99 assay marks therefore encode 52 distinct reporter-level values; assays for the same reporter have identical cell scores. Reporter-level comparisons use two-sided paired Wilcoxon signed-rank tests against the prespecified MLP, Holm correction across 54 comparisons and an absolute median paired-effect threshold of 0.02. Screen and assay summaries are descriptive.

Eleven of 54 paired comparisons against the prespecified reporter-specific multilayer perceptron (MLP) met Holm-adjusted *P <* 0.05 and a median paired-effect threshold of 0.02. All favoured the MLP over Ridge, scButterfly or MIDAS in field or gene tests (Fig. 2b). Held-out-gene median recovery was 0.768 for MLP, 0.691 for scButterfly and 0.614 for MIDAS; within-reporter MLP gains were 0.081 and 0.133 (*n* = 52, both adjusted *P <* 2 *×* 10^*−*8^). The gain over Ridge was 0.018, below the effect criterion despite statistical significance. Other comparisons were descriptive.

Descriptive rankings also depended on the test. The MLP had the highest reporter-level knockout-response median with held-out fields (0.715), whereas Ridge had the highest whole-screen median (0.730). The latter exceeded the MLP median of 0.677, but the median paired Ridge-minus-MLP difference was *−* 0.002 across the same 34 reporters (Holm-adjusted *P* = 1). Target-matched comparisons thus gave a different view from the ordering of pooled medians. Screen and assay views localized variation across the acquisition topology (Fig. 2c,d); field/gene cell-level panels reweighted reporter scores by observed membership rather than supplying additional independent cell-level estimates.

The recoverability landscape was summarized by the unweighted median of method *scores*, with screen and assay distributions showing variation across biological systems (Supplementary Figure 5a). For fidelity and response replacement, we instead evaluated a fixed ensemble: the element-wise median of aligned out-of-fold *predictions*. This separates a summary of the available predictors from assessment of a particular inferred readout.

### Response diagnosis and use decisions provide complementary assessments

Median method scores varied widely across reporters (Fig. 3a). Eight of 52 reporters had held-out-gene recovery below 0.60; this rose to 12 of 34 repeated reporters in screen transfer. Mean gene-test recovery ranged from 0.877 for endosomal reporters to 0.620 for signalling reporters. Intensity and intensity-range endpoints were more recoverable (0.822 and 0.791) than spatial displacement and sub-object distribution (0.541 and 0.525; Fig. 3b).

**Figure 3.**
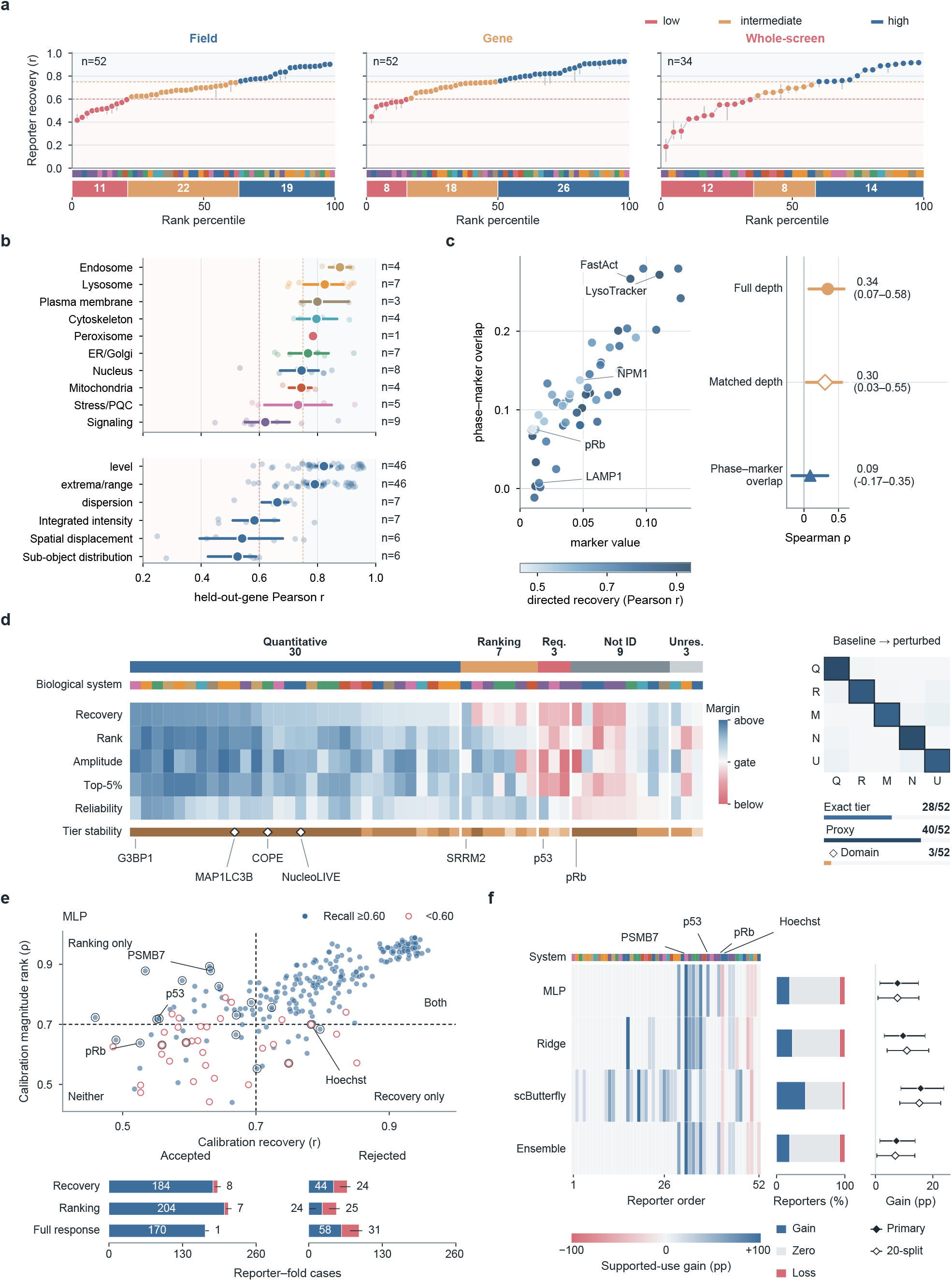
Response diagnosis and calibration-based use decisions. **a**, Reporter recoverability across field, gene and screen tests (*n* = 52, 52, 34), summarized by median method scores; bands denote descriptive strata, not sufficiency tiers. **b**, Recoverability by biological system and endpoint family; points are reporter-level values, means and reporter-bootstrap 95% intervals (10,000 resamples). **c**, Perturbation distinctiveness and phase–reporter overlap versus held-out-gene recovery (*n* = 52); intervals use 3,000 reporter resamples. Matched-budget values use one draw capped at 300 cells per sgRNA. **d**, Five criterion margins and fixed ensemble tiers for 52 reporters; insets show threshold sensitivity and domain flags. **e**, Calibration recovery and magnitude rank for 260 MLP reporter–fold cases; colours encode four evaluation outcomes and bars compare acceptance rules. Evaluation success requires top-20% hit recall *≥* 0.60. **f**, Rank-minus-recovery gain in supported successes, as percentage of all cases, for four predictors. The matrix retains every reporter; right summaries compare the primary partition and mean of 20 fixed partitions, with 2,000 reporter-cluster bootstrap intervals.

We compared how strongly a reporter distinguishes perturbations, how similar its perturbation structure is to phase, and whether phase predicts its response to an unseen knockout perturbation (Fig. 3c). Reporter perturbation distinctiveness was modestly associated with gene-test recovery (*ρ* = 0.344, 95% interval 0.072–0.579, *n* = 52); one subsampling draw capped at 300 cells per guide gave *ρ* = 0.305. Phase–reporter overlap was nearly unrelated to recoverability (*ρ* = 0.090, *−* 0.174 to 0.345). Similarity between perturbation-distinctiveness profiles therefore provided little guidance to prediction performance on unseen perturbations. These quantities describe different aspects of a reporter’s value for a morphological screen.

Atlas-specific rules classified the fixed ensemble into 30 quantitative proxies, 7 ranking proxies, 3 measurement-required, 9 not-identifiable and 3 unresolved assignments (Fig. 3d). Quantitative proxies met specified correlation, magnitude-spread and hit criteria, rather than demonstrating per-endpoint calibration. Not identifiable denoted low or unavailable reference reliability; unresolved included missing or intermediate fidelity evidence. Across 36 threshold variations, 40 reporters retained proxy/nonproxy status and 28 retained their exact tier (Supplementary Table 5).

We next divided each MLP outer-test fold into calibration and evaluation genes without refitting the predictor, producing 260 reporter–fold cases. The evaluation task selected the strongest 20% responses and required hit recall *≥* 0.60 (Fig. 3e). Accepting every case would yield 228 successes and 32 failures, a failure rate of 12.3%. Recovery-only accepted 192 cases: 184 supported successes and 8 supported failures, with 44 missed successes and 24 true rejections. Rank-only accepted 211: 204 successes and 7 failures, with 24 missed successes and 25 true rejections. Full-response accepted 171: 170 successes and 1 failure, with 58 missed successes and 31 true rejections. Full-response combined recovery, rank, variance and calibration hit evidence; it did not include a repeat gate in this experiment.

Thus, rank-only supported 20 more successful cases than recovery-only, with one fewer supported failure. Rank-minus-recovery supported-use gain was 7.69 percentage points (reporter-cluster 95% interval 1.54–15.00); gains were also positive for Ridge, scButterfly and the ensemble (Fig. 3f). Full-response offered a more selective operating point, reducing false support from 3.3% to 0.6% of accepted cases compared with rank-only, while reducing coverage from 81.2% to 65.8%. At matched 75% coverage, the MLP false-support difference was *−*2.05 points (*−*5.85 to 1.14), compatible with no difference (Supplementary Figure 6; Supplementary Data 1). Ranking evidence therefore expanded the set of demonstrated prioritization uses; combined response criteria selected a smaller set with fewer failures. The appropriate operating point depends on how many missed opportunities can be accepted in exchange for fewer unsuccessful uses.

Keeping reliability and thresholds fixed, 20 of 43 reproducible reporters were proxies under every predictor, 6 under none and 17 crossed the proxy boundary. Fifteen consistently supported reporters changed between quantitative and ranking tiers. LAMP1 was quantitative under MLP, ranking under MIDAS and nonproxy under scButterfly (Supplementary Figure 3). Predictor choice changed supported uses as well as correlation.

Paired-data availability and representation also affected recovery (Supplementary Figure 2). In a 12-reporter label-efficiency analysis, mean recovery improved from 0.1% to 1% of available paired measurements for all ten methods and further at 20% for nine. Across six reporters, mean gene-test recovery was 0.739 with engineered features, 0.594 with frozen Cytoland and 0.613 after partial fine-tuning. Fine-tuned Cytoland remained below engineered features (paired difference *−*0.127, reporter-hierarchical 95% interval *−*0.188 to *−*0.066). These comparisons show how representation and paired-data availability affect recovery, one component of the proxy assessment.

### Same-cell morphology contributes information beyond perturbation averages

Prediction could reflect differences between gene–screen groups even if phase and reporter phenotypes were unrelated among cells within each group. We fixed the MLP and altered only pairing or phase-feature content for 52 reporters across five gene holdouts (Fig. 4a). Within-gene–screen target reassignment preserved response distributions but broke same-cell linkage during fitting; test targets retained exact pairing. Exact pairing gave median cell/knockout correlations of 0.757/0.769, compared with 0.291/0.293 after reassignment. Reassigning targets among cells matched for area, density and eccentricity gave 0.531/0.502, and prediction using size–shape features alone gave 0.261/0.329.

**Figure 4.**
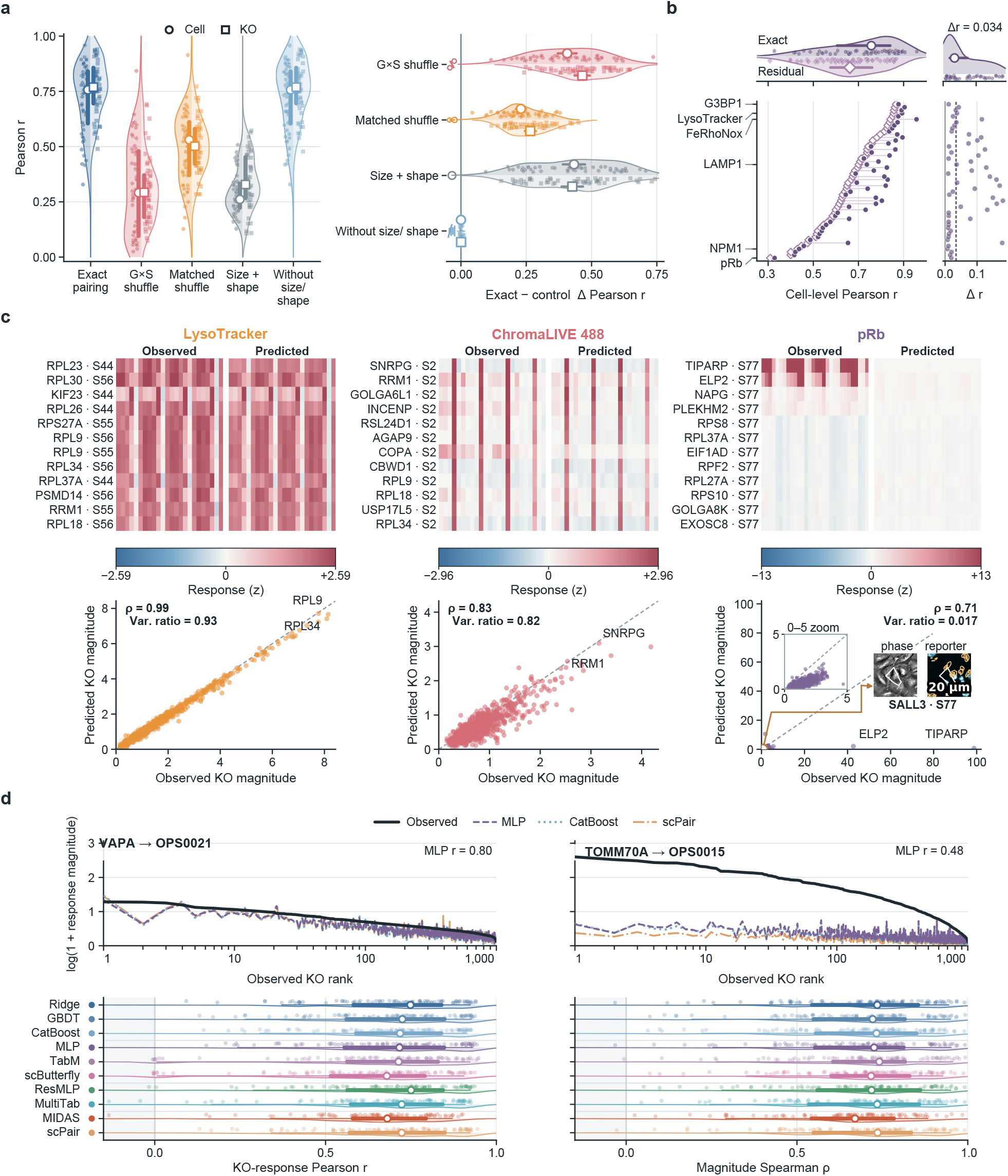
Targeted-state fidelity has separable cell, perturbation and transfer limits. **a**, Reporter-specific MLP recovery under exact pairing and four information interventions (*n* = 52, five held-out-gene partitions); G*×* S denotes gene–screen shuffling and matched denotes covariate-matched shuffling. Circles/squares denote cell/knockout scores, pale lines pair reporters, and open effect symbols show means with reporter-bootstrap 95% intervals. **b**, Exact-pair prediction versus an MLP fitted to cross-fitted within-gene–screen target residuals (*n* = 52), with ranked differences. Residuals use targeted measurements from peer cells in the held-out condition. **c**, Observed versus median-of-ten-method predictions on a common reporter-wide control reference: 12 strongest held-out gene–screen profiles per heatmap and all 1,000 genes per full-range scatterplot; the pRb zoom shows 0–5 on both axes. Screen labels use the source OPS numbering; S77 denotes OPS0077. The microscopy inset identifies an exact SALL3 cell from OPS0077 (S77); scale bar, 20 *µ*m. **d**, Transfer curves show two prespecified reporter-to-destination directions and three representative predictors. Half-rainclouds retain all 81 directed transfers per method, displaying density, median, interquartile range and 5th–95th percentiles for response reconstruction and magnitude ranking.

Removing size and shape left performance essentially unchanged. Same-cell morphology therefore carried information beyond group averages and these measured covariates.

Cross-fitted gene–screen residual targets further isolated within-condition variation (Fig. 4b). Each cell’s residual used a condition mean estimated from other cells, excluding its own evaluation block. Residual correlations were positive for all 52 reporters (median 0.662), compared with 0.757 for exact-cell correlations. The residual reference uses target measurements from peer cells in the held-out condition, so this analysis evaluates conditional within-condition association. Together, the pairing and residual controls show information about individual-cell targeted states beyond the average response of a perturbation group.

Individual-cell information did not ensure faithful perturbation magnitudes. Ensemble predictions gave magnitude rank/variance ratios of 0.988/0.930 for LysoTracker, 0.828/0.816 for Chroma-LIVE 488 and 0.708/0.017 for pRb (Fig. 4c). Across 810 predictor–transfer combinations, median response Pearson was 0.715 and magnitude Spearman 0.723, with marked variation between destinations (Fig. 4d). Ranking and response spread were distinct capabilities; pRb additionally lacked a reproducible reference.

### Reference stability and response contraction constrain different uses

Gene-test recovery correlated with within-screen reliability (*ρ* = 0.547, 95% interval 0.295–0.737), cross-screen consistency (0.349, 0.014–0.625) and coverage (0.482, 0.211–0.686; Fig. 5a). Guide consistency had a similar direction but an interval including zero (0.376, *−* 0.002 to 0.668). Reliability, guide consistency and log coverage jointly accounted for 38.7% of in-sample recovery variation among 35 complete cases (permutation *P* = 0.001).

**Figure 5.**
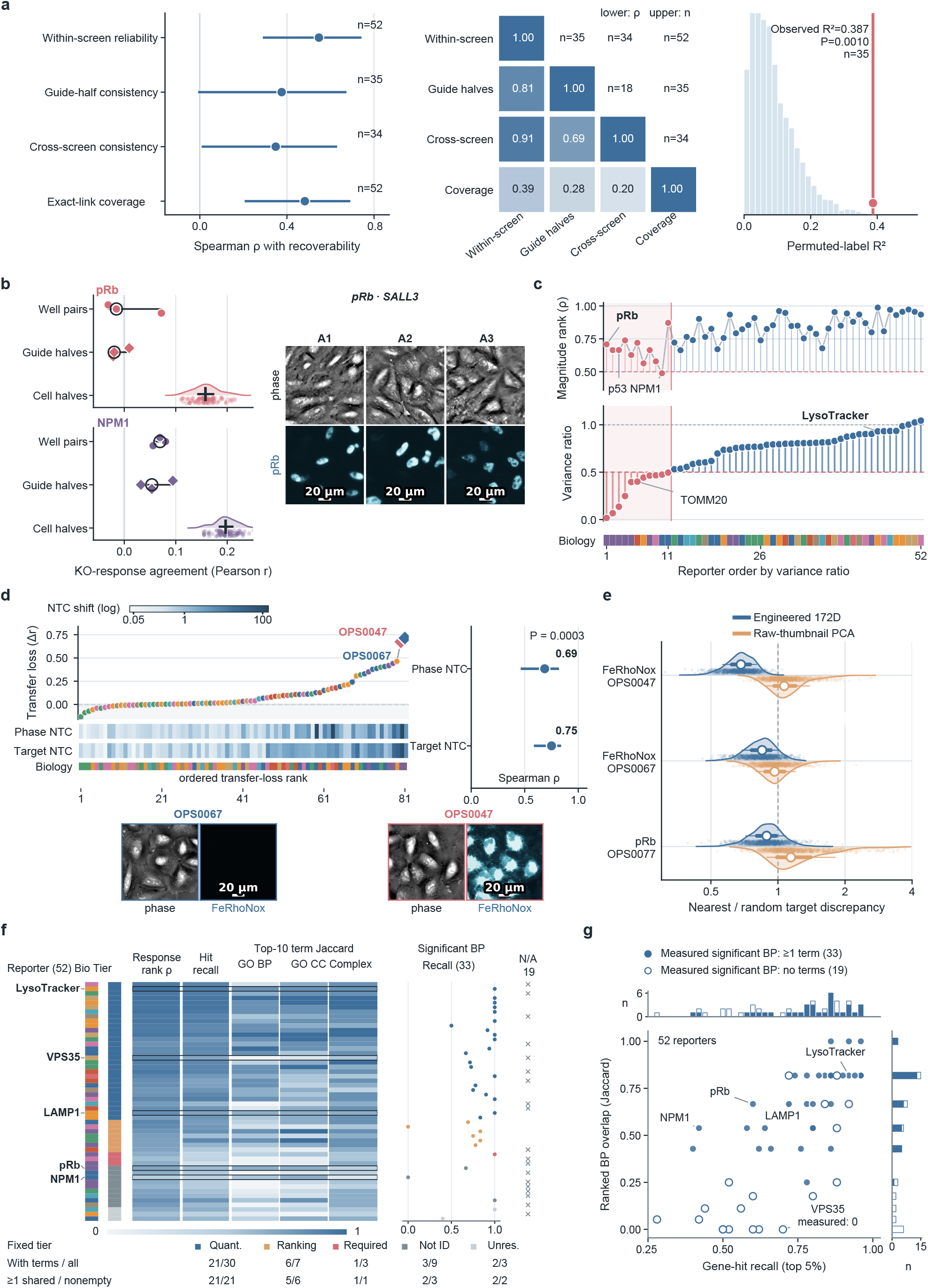
Reference stability, response fidelity and retained functional evidence. **a**, Recoverability versus reference consistency and coverage; reporter-bootstrap 95% intervals. **b**, pRb and NPM1 across three well pairs, three guide splits and 100 random cell splits. **c**, Magnitude rank and variance ratio for 52 reporters; red marks ratios below 0.5. **d**, Control-cell shifts versus transfer loss: 81 directions (displayed correlations), 34 reporter medians (inference). Diamonds mark FeRhoNox examples. **e**, Nearest/reference target discrepancy for engineered features and raw-phase principal components; bars, medians and interquartile ranges. Scale bars, 20 *µ*m. **f**, Fixed reporter order and tiers with response, hit and top-ten term retention; right tracks show significant biological-process retention at BH-adjusted *P≤* 0.05. Recall is undefined for 19 empty measured sets. **g**, Hit recall versus ranked biological-process overlap for 52 reporters (45 distinct coordinates); filled/open points denote nonempty/empty measured significant sets. Marginals retain exact counts without jitter.

Median agreement across well pairs, guide halves and random cell halves was *−* 0.015, *−* 0.020and 0.158 for pRb, and 0.069, 0.054 and 0.197 for NPM1 (Fig. 5b). Random-cell splits evaluated sampling stability within the observed population; well and guide comparisons included variation across acquisition units and perturbation reagents. Agreement across wells and guide subsets was lower than agreement across random cell halves. For these reporters, improving the reproducibility of the reference response is a prerequisite for distinguishing model error from variation in the measured perturbation effect.

Contraction of response magnitudes represented a different constraint. Across 52 reporters, median magnitude rank was 0.825, variance ratio 0.779 and top-5% recall 0.790. Eleven reporters retained less than half the measured magnitude variance; ten still had rank correlations above 0.5 (Fig. 5c). These reporters preserved much of the perturbation hierarchy while compressing the differences in response magnitude between genes. Such predictions can remain useful for selecting candidates even when comparing effect sizes requires more accurate amplitude calibration.

Control-cell shifts were associated with transfer loss across 81 directions; reporter-level correlations were 0.765 for phase and 0.837 for targets (both permutation *P* = 1/3,001; Fig. 5d). The control cells carry no targeted knockout, making their shifts an indicator of changed acquisition or cellular context. In the displayed FeRhoNox examples, changes in control-cell appearance accompanied deterioration in prediction.

Morphology-nearest cells also retained substantial target discrepancy, with nearest/reference ratios of 0.681 and 0.849 in two FeRhoNox contexts and 0.889 for pRb (Fig. 5e). A ratio near one means that selecting a morphologically similar neighbour removes little of the discrepancy in targeted state. Raw-thumbnail comparisons gave ratios near or above one. These results identify residual target variation within the measured representation, which can contribute to averaging and response contraction. Reference instability, acquisition shifts and unresolved cellular variation consequently suggest different priorities for improving the measurement.

### Response replacement separates hit retention from functional evidence

We replaced observed responses to held-out knockouts with ensemble predictions and repeated the same downstream analyses. The first task prioritized the 50 strongest perturbation responses; the second compared functional annotations associated with these hits, using Gene Ontology biological processes and cellular components ^31,32^ and curated protein complexes ^33^. Predictions retained a median 79% of the strongest hits. Median Jaccard overlap between the ten leading terms was 0.603 for biological processes, 0.667 for cellular components and 0.667 for protein complexes.

We then evaluated significant terms separately against the 1,000 assayed genes, using BH-adjusted *P ≤* 0.05 (Fig. 5f,g). Thirty-three reporters had nonempty measured biological-process sets; 19 did not. Among quantitative proxies, 21 of 30 had such evidence and all 21 retained at least one significant term. Significant recall was undefined for empty measured sets. This separates the stability of a fixed-length ranked list from retention of biological associations supported by the measured screen.

VPS35 illustrates why ranked lists and significant evidence must be distinguished. Its top-ten biological-process lists had zero overlap at the 5% hit budget, but neither list contained a significant term. At 10% and 20% budgets, predictions retained four of five and five of eight measured significant terms, respectively (Supplementary Data 1). The six reporters with zero top-ten overlap all lacked comparable measured significant evidence at the primary budget. Ranked-list mismatch thus did not establish loss of a significant biological programme.

An auxiliary cohort-relative downstream-retention index combined strong-hit recall and ranked-term overlap (Supplementary Figure 5b,c). A nested leave-one-reporter-out Ridge analysis related fidelity and reliability to this index (*ρ* = 0.786, *R*^2^ = 0.587; Supplementary Figure 4a). It summarized variation in retained analyses across reporters, whereas the task outcomes directly measured performance against a specified selection criterion.

### Independent assay resources test task-specific uses and decision transfer

External applications changed biological context, readout and pairing unit (Fig. 6). A resource survey recorded 59 candidates, with 42 assessable for eligibility (Supplementary Table 1); harmonized collections also informed this survey ^34^. OASIS paired brightfield and biochemical readouts in primary hepatocytes ^35^; Cell Health paired fluorescent morphology and separately acquired health end-points ^15^; LINCS paired morphology and RNA at matched compound conditions. PERISCOPE supplied a fluorescent-channel reference ^10^.

In primary hepatocytes, we asked whether brightfield predictions could prioritize compounds causing the strongest metabolic-activity losses for follow-up measurement. Using released bright-field DINO features, Ridge was fitted on 868 development compounds and tested on 217 held-out compounds, grouping every dose and source. The ranking score was the signed mean loss of RealTime-Glo metabolic-activity signal relative to same-plate DMSO, averaged over the assayed concentrations. Selecting the predicted top 5% recovered 10 of the 11 largest measured losses (90.9%; compound-bootstrap interval 54.5–100%; Fig. 6a,b). Mean measured loss was 0.334 in the predicted selection and 0.338 in the measured top eleven, showing retention of substantial metabolic decreases. Among 94 compounds with complete dose-matched measurements in two production sources, predictions from one source recovered four of the five strongest measured losses in the other, in both directions; measured-repeat selection also recovered four of five. These source comparisons supplied a repeat reference with wide uncertainty (Supplementary Figure 4c). Histogram gradient boosting (HGB) also recovered 10/11. Matched predictors using fluorescence-derived cell counts recovered 8/11 with Ridge and 7/11 with HGB; acquisition metadata alone recovered 0/11 and 1/11, respectively (Supplementary Data 3). Disrupting within-plate prediction–compound correspondence reduced mean brightfield-Ridge recall to 5.38%. Fixed 10% and 20% budgets retained 17/22 and 34/44 hits. These results support prioritization of strong metabolic losses at several follow-up budgets.

**Figure 6.**
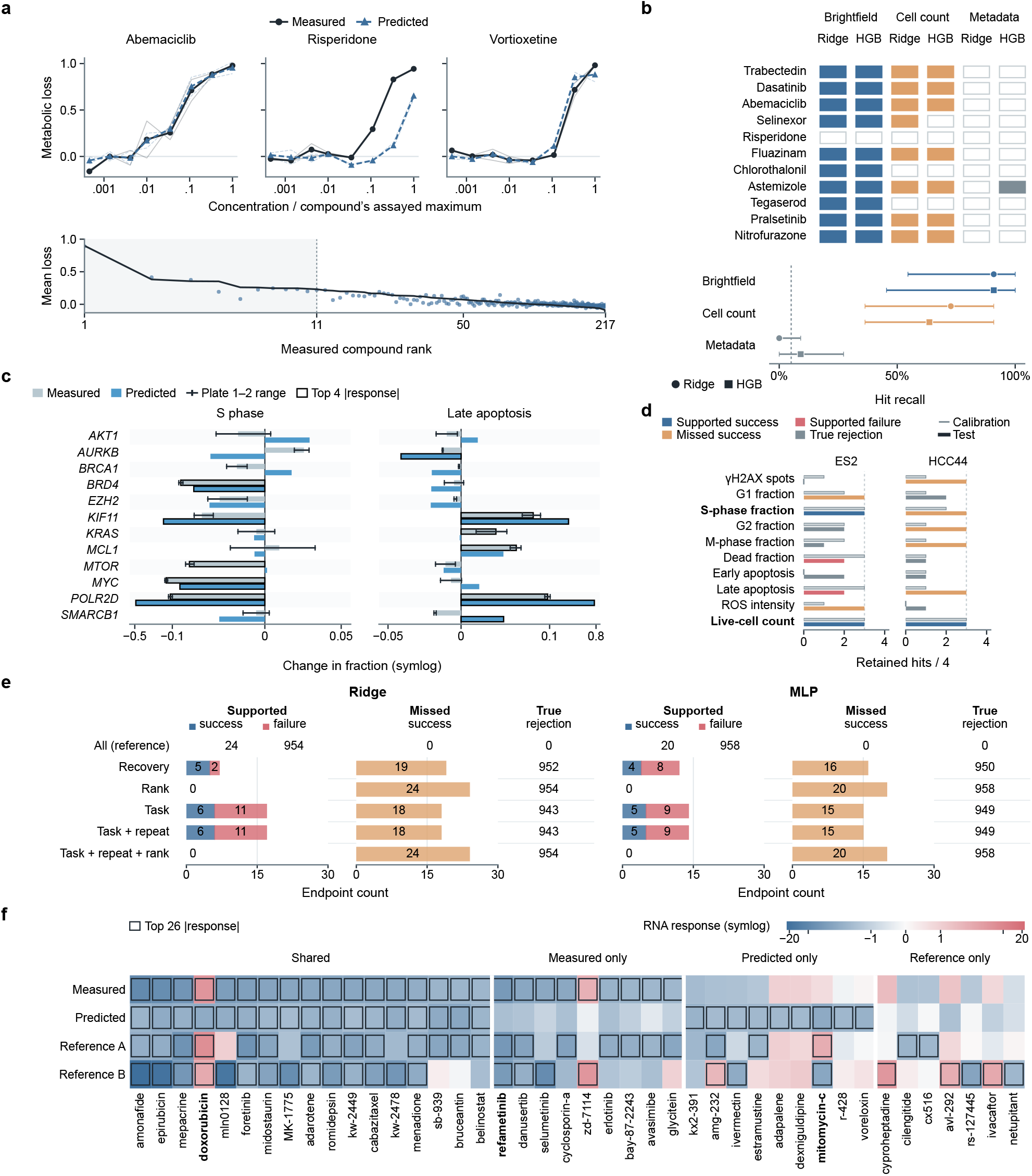
Independent resources connect response prediction to specific measurement uses. **a**, OASIS primary-hepatocyte examples: measured and brightfield-Ridge-predicted metabolic-activity loss across the deposited dose range for three compounds, followed by the complete 217-compound held-out ranking. The dose coordinate is each compound’s assayed concentration divided by its assayed maximum; the loss is the signed change from same-plate DMSO. **b**, Recovery of the eleven strongest measured losses by Ridge and histogram gradient boosting (HGB) from brightfield, fluorescence-derived cell-count and acquisition-metadata inputs. Filled cells are recovered hits; lower intervals are compound-bootstrap 95% intervals, with the fixed random-selection expectation shown. **c**, Cell Health responses for twelve held-out genes in ES2: measured consensus (grey bars), predictions (blue bars), two-plate ranges (whiskers) and the four largest absolute responses in each set (black outlines). S-phase and late-apoptosis fractions use symmetric-log axes, linear within *±* 0.01. **d**, Cell Health task outcomes across twenty prespecified endpoint–context cases. Calibration and test retained-hit counts are shown separately out of four; colours distinguish supported success, supported failure, missed success and true rejection. **e**, LINCS endpoint decisions for Ridge and MLP across five gated rules plus an ungated reference. Supported successes and failures share a 0–30 count scale with missed successes; true rejections are labelled directly. **f**, LINCS BRCA1 response map for all 42 compounds in the union of measured, predicted and two separately acquired reference top-26 sets. Colours show signed, plate-DMSO-centred RNA responses on one symmetric-log scale; outlines mark top-26 membership.

The broader OASIS CellProfiler assessment asked how well the complete biochemical response was reconstructed. It gave metabolic-activity and lactate dehydrogenase (LDH) release recovery of 0.83 and 0.67, but magnitude ranks of 0.604 and 0.633. Both were unresolved under the full response tiers despite top-5% recall of 0.727 (Supplementary Table 2). The focused analysis differs in representation, signed score and aggregation, and directly evaluates the intended selection. Thus, incomplete preservation of the broader response ordering can coexist with a useful prioritization task in the same resource.

Cell Health used ten endpoints in ES2 and HCC44, giving 20 cases, with 25 training, 12 calibration and 12 evaluation genes. In ES2, S-phase prediction retained three of four strongest responses at *r* = 0.604, whereas late-apoptosis prediction retained two of four at *r* = 0.950 (Fig. 6c). The calibration task rule accepted five cases: three supported successes and two failures, with seven missed successes and eight true rejections (Fig. 6d). Adding plate-repeat agreement left the primary acceptance set unchanged. HCC44 live-cell count passed against the plate consensus (3/4), but not the second plate (2/4). Random selection has expected recall 1/3 at this four-of-twelve budget, and each retained gene changes recall by 0.25.

LINCS extended the assessment to RNA, evaluating all 978 landmark probes against 260 held-out compounds. BRCA1 predictions retained 17 of the 26 strongest absolute responses against the consensus and each of two separately acquired RNA references, at signed *r* = 0.515 (Fig. 6f). Sixteen of the seventeen consensus-recovered hits agreed in response direction; thirteen recovered hits were common to both reference comparisons. This supplied an RNA prioritization example at moderate overall correlation, complementing the Cell Health example in which high correlation did not preserve enough strong-response hits.

We also tested whether calibration evidence could select successful RNA uses across the full probe panel (Fig. 6e). Recovery-only accepted seven Ridge cases (five successes, two failures), whereas task-only accepted seventeen (six successes, eleven failures). MLP accepted twelve recovery cases (four successes, eight failures) and fourteen task cases (five successes, nine failures). Task-only success fractions were 35.3% and 35.7%, with coverage of 1.7% and 1.4%; the same rules missed 18 and 15 successful endpoints. Repeat gating preserved the primary acceptance sets, and rank gating accepted none. Across ten Ridge partitions, the task-plus-repeat failure fraction ranged from 40.0% to 66.7%. The panel-wide analysis therefore found useful RNA tasks but limited consistency in identifying them from calibration evidence. It measures a different requirement from demonstrating hit retrieval for an individual target.

PERISCOPE recovered anti-TOMM20 responses from four fluorescent predictor channels but had guide-partition reliability of 0.005 and 0.047 at cell and guide-profile levels. Core-essential and mitochondrial subsets also remained below 0.30. Input-permutation and leave-one-dye-out controls separated prediction failure from insufficient reference reproducibility (Supplementary Figure 1). Across the external resources, the same assessment connected predictive performance to a concrete task and identified which additional evidence was needed to extend that use.

## Discussion

Whether a predicted readout could replace a measurement depended on the analysis it had to support. In a paired perturbation atlas, label-free morphology carried information about fluorescently measured cell states beyond perturbation averages and measured covariates. Predictions often preserved the ranking of knockout effects while compressing their magnitudes. Such predictions can support the prioritization of strong perturbations even when comparing effect sizes requires better quantitative fidelity. Beyond the atlas, high correlation did not always retain the strongest responses in cell-health data, whereas moderate correlation sufficed for an RNA target. Evaluating the intended analysis directly connects the promise of virtual-cell models to an experimentally testable question: which biological analyses can an inferred readout support?

The pairing controls help explain what morphology contributes. Reassigning targets among cells of the same perturbation and screen preserved group averages but reduced median cell-level recovery from 0.76 to 0.29. The additional predictive information therefore depended on individual-cell linkage, beyond perturbation means. Yet morphologically similar cells retained substantial differences in targeted state. This unresolved variation offers one explanation for response contraction: when a predictor estimates the average target state compatible with its input, it can narrow the spread of responses. The observed coexistence of retained rankings and compressed magnitudes is consistent with this averaging.

Method choice is consequential within this framework. Paired reporter comparisons supported MLP gains over several comparators, while recomputing response-fidelity tiers for each predictor showed which gains changed the supported use. Some targets remained proxies across all assessed predictors, whereas others changed their quantitative, ranking or proxy-use status. Model improvement thus has an interpretable outcome: a better predictor can enable a measurement use that was previously unsupported. The 30 quantitative and 7 ranking proxies describe the assessed ensemble in its acquisition and training context; subsequent predictors can be evaluated against the same biological tasks.

Calibration and evaluation separated the choice of a predicted readout from the test of its biological use. In OPS, ranking-based acceptance supported 204 successful prioritization cases compared with 184 for recovery-only, with seven versus eight supported failures. Combining response criteria supplied a more selective set with one supported failure. These operating points serve different experimental priorities: broad prioritization benefits from retaining more usable targets, whereas costly follow-up can favour fewer unsupported selections. The matched-coverage comparison had uncertainty spanning no difference, so the clearest demonstrated gain was in successful-use coverage. Reporting both coverage and failures makes this choice visible to the experimenter.

The external applications broadened the assessment beyond phase images and same-cell fluorescence. Hepatocyte brightfield profiles selected ten of eleven strong metabolic-activity losses, compared with eight for a matched Ridge predictor using fluorescence-derived cell counts. Repeated biochemical measurements supplied a reference for the same follow-up selection. Cell Health and LINCS showed from opposite directions why predictive correlation alone can misrepresent a use: high correlation did not always retain the required strong hits, while BRCA1 retrieval succeeded at moderate correlation. Across the complete LINCS probe panel, calibration task performance often changed on evaluation compounds. A successful target-specific application and reliable selection of such applications across a panel are distinct outcomes; the calibration–evaluation analysis tests the latter directly.

Reference reproducibility determines how well model error can be distinguished from variation in the measured response. Nine reporters fell below the atlas reliability criterion, limiting the evidence for response replacement even when some predictive metrics were high. Functional analyses likewise require an interpretable measured reference. Twenty-one quantitative proxies with significant measured biological-process evidence each retained at least one significant term. VPS35’s zero ranked-term overlap occurred when neither list was significant at the primary budget, and increasing the fixed hit budget revealed shared significant terms. Ranked-list agreement describes stability among the leading annotations; significant-term recall describes retention of associations supported by the observed screen. Evaluating both clarifies how hit selection and available reference evidence shape the biological interpretation.

Practical application starts with a paired pilot and a holdout matching the intended deployment unit. The task and tolerance determine which properties must be retained, and relevant repeat measurements establish the available reference. Cell-half, guide and well comparisons capture different combinations of sampling, reagent and acquisition variation; their choice follows the proposed use. A predictor can then be fitted, acceptance fixed using calibration perturbations and the intended analysis repeated on evaluation perturbations. Low repeat agreement points towards improving the reference; sensitivity to screen context points towards validation in the intended acquisition setting; residual ambiguity within morphology motivates richer representations or additional measurements. The atlas conclusions were established in A549 cells using a 172-dimensional engineered morphology representation and paired training data. The six-reporter Cytoland comparison favoured engineered features, while further learned representations offer a way to test whether more spatial information can be recovered ^36–38^. The 12-reporter label-efficiency analysis similarly shows that the paired-data budget is part of the measurement design (Supplementary Figure 2). MorphoSuff accepts paired measurements or supplied predictions, allowing these choices to be evaluated with the same evidence and task outcomes across datasets. Measurement sufficiency is therefore a property of a target, predictor, representation and context, assessed for a particular analysis. It lets experimenters use paired pilot data to decide which readouts a screen can infer and which it must still measure.

## Methods

### Study design, data resources and phenotype representations

We analysed the public optical perturbation atlas of A549 cells ^27^; no new biological samples were generated. The source study combines pooled CRISPR knockouts, phase and targeted-fluorescence imaging, perturbation identities obtained by in situ sequencing and engineered single-cell phenotypes. The analysed data comprised 1,000 knockout genes, 52 reporters, 73 physical screens and 99 observed reporter–screen assays spanning ten biological systems. Quality-controlled linkage yielded 7,344,374 distinct phase cells and 9,996,286 exact-linked phase–reporter observations because one phase cell could contribute more than one reporter observation. Reporter–screen combinations that were not acquired were treated as missing assays, never as negative measurements.

Reporter was the primary inferential unit for atlas-wide claims. Physical screen and reporter– screen assay were additional reporting units; genes, endpoints, cells and data partitions were nested observations. Dependence among directed whole-screen tests was handled using reporter-level summaries and reporter-preserving resampling, as specified below. Inclusion required a valid source linkage, an observed reporter endpoint block and assignment to a prespecified partition. Exclusions and unresolved mappings are reported in the deposited panel-level source tables.

### Phenotype representations

The primary input was a 172-dimensional engineered morphology vector derived from each segmented phase image. The features described image intensity, texture, object geometry and spatial organization. The target was a reporter-specific engineered phenotype derived from the fluorescence image of the same cell. The target schema comprised one 20dimensional block, 38 24-dimensional blocks, six 30-dimensional blocks, one 60-dimensional block and six 72-dimensional blocks, for 1,604 endpoints in union. Fitting and evaluation used only the endpoints observed for a reporter; heterogeneous blocks were not padded with synthetic targets.

Model preprocessing was fitted within the training portion of each partition. Phase variables finite in fewer than 80% of training cells were removed; remaining non-finite values were replaced by the training median, variables with training standard deviation no greater than 10^*−*8^ were removed, and retained variables were centred and scaled by their training means and standard deviations. A cell was excluded for a reporter if any endpoint in that reporter’s observed target block was nonfinite. Target endpoints were standardized by reporter using training means and standard deviations, with invariant endpoints removed. The fitted feature filters, imputation values and scaling parameters were applied unchanged to validation and test cells. Size and shape features were identified from the feature schema before the same-cell controls were evaluated. Cell, gene, guide, field, screen and reporter identifiers were retained in every prediction table to reconstruct the evaluation units independently of phenotype values.

### Prediction models and generalization framework

Three prespecified evaluation regimes represented distinct deployment questions. For held-out fields, the physical screen–well–tile identifier was assigned once in the global phase table and hashed into five folds. The same physical field, and any exact-linked phase row reused by different reporter heads, therefore had one fold assignment throughout specialist and shared-model training; knockout identities and screen context remained represented in training. For held-out genes, knockout genes were partitioned into five folds and every cell carrying a test gene was excluded from model fitting and model selection. This test evaluates response reconstruction for an unseen perturbation within the observed reporter–screen structure. In both fivefold analyses, fold *k* was the outer test set, fold (*k* + 1) mod 5 was the validation set and the other three folds were used for training.

For strict whole-screen transfer, all observations from one physical destination screen were removed from training. A valid direction required the same reporter in at least one source screen and in the held-out destination. This produced 81 directed evaluations across 34 repeated reporters and 66 destination screens. Held-out-field and held-out-gene summaries included 52 reporters, 73 physical screens and 99 reporter–screen assays; whole-screen summaries included 34 reporters, 66 destination screens and 81 directed assays. Reporter scores were arithmetic means over the five partitions for field and gene tests and over eligible destination-screen directions for whole-screen transfer.

### Prediction methods

The ten methods were selected before the final comparative analysis: Ridge regression, gradient-boosted decision trees, CatBoost ^39^, a reporter-specific MLP, TabM ^40^, scButterfly ^41^, a shared residual MLP, MultiTab, MIDAS ^42^ and scPair ^43^. The set covers conventional target-specific tabular prediction, neural tabular models, shared multitask prediction and single-cell multimodal completion. Every method received the same 172-dimensional input, observed reporter-specific endpoints and train, validation and test partitions; training scope, loss functions and model selection are summarized in Supplementary Table 6.

Target-specific methods were fitted separately for each reporter. Shared methods used a common representation with reporter-aware outputs or endpoint masks, and their loss was averaged over observed endpoints only. Hyperparameters and stopping rules were selected on development data and fixed before final evaluation. All downstream analyses used out-of-fold predictions. The benchmark used the validation-selected single-model states; its recoverability landscapes were summarized by the unweighted median of ten method-level scores. The response-replacement ensemble instead combined predictions from the archived parameter-averaged checkpoints of TabM, scButterfly, ResMLP, MultiTab, MIDAS and scPair, with the same fitted Ridge, GBDT, CatBoost and MLP predictors as in the benchmark. Each averaged checkpoint supplied one model prediction; the method-specific states and input files are identified in Supplementary Data 1. For each reporter, fold and gene–screen row, we took the element-wise unweighted median of the ten aligned response vectors before averaging across screens. Held-out-gene fidelity, measurement-sufficiency classification and utility-prediction analyses evaluated this fixed ensemble; their recoverability was calculated directly from its predictions using Eq. 2, followed by the arithmetic mean over five folds. No method received performance-dependent weighting, and methods were not treated as biological replicates.

Published comparator implementations were adapted to accept the common phase input, heterogeneous reporter blocks and predefined partitions. Unobserved reporter endpoints were masked from both the loss numerator and denominator rather than encoded as zeros. Model selection used validation outcomes only. Supplementary Table 6 specifies the architecture, objective, training scope and selection procedure used for each implementation.

### Recoverability metrics and aggregation units

Let *I*_*rsg*_ denote exact-linked cells for reporter *r*, screen *s* and knockout gene *g*, and let *I*_*rs*0_ denote non-targeting controls from the same reporter and screen. For endpoint *e*, the control-relative response was

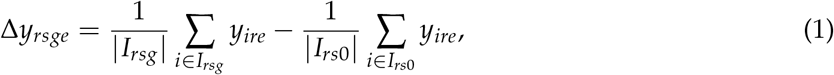

with Δ*ŷ*_*rsge*_ defined analogously from out-of-fold predictions. Cell-level recoverability was calculated separately for every observed endpoint across exact-linked test cells and then averaged equally across endpoints. Knockout-level recoverability was the corresponding macro-endpoint statistic,

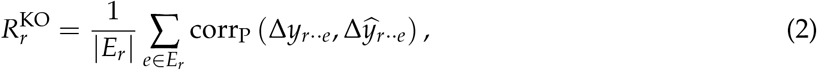

where *E*_*r*_ is the observed endpoint set for reporter *r* and the correlation runs over eligible screen–gene response rows in the relevant test partition. A Pearson value required at least two paired observations and non-zero observed and predicted variance; otherwise that endpoint was undefined and excluded from the equal-endpoint macro-average. A screen–gene response could contain one or more targeting cells but required matched test controls from the same screen. Non-targeting controls define Eq. 1 but are excluded from the cell-level correlation. Scores were then summarized by reporter, physical screen or reporter–screen assay, corresponding respectively to target recoverability, environmental recoverability and recoverability of a particular target in a particular environment. For each method, knockout-response scores for individual reporter–screen assays were computed within each outer partition and averaged over the five field or gene partitions; physical-screen scores were equal-weight means over member assays. The same aggregation was used in the benchmark and recoverability landscape. For held-out-field and held-out-gene cell-level screen and assay views, common per-cell predictions were unavailable for every shared method; these views therefore reweighted reporter-level cell macro-Pearson values by observed screen or assay membership, with equal assay weights within screen. Strict whole-screen cell scores were evaluated directly in destination cells.

Reporter-level method comparisons used a prespecified reporter-specific MLP reference and two-sided paired Wilcoxon signed-rank tests over reporters. A difference was called statistically and practically distinct only when the Holm-adjusted *P <* 0.05 across 54 comparisons and the absolute median paired effect was at least 0.02. Tests paired 52 reporters in the field and gene views and 34 repeated reporters in whole-screen transfer. Screen and assay summaries were descriptive because their units share reporters, genes and physical acquisition structure. Descriptive low-, intermediate- and high-recoverability strata were defined as Pearson *r <* 0.60, 0.60*≤ r <* 0.75 and *r≥* 0.75, respectively. These strata summarize the recoverability distribution and were not used as measurement-sufficiency tiers.

### Biological and phenotype summaries

Reporters were assigned to ten biological systems using source metadata. Endpoints were assigned to six families: intensity level, intensity extrema or range, intensity dispersion, integrated intensity, sub-object distribution and spatial displacement. Biological-system means used reporter as the unit. Endpoint-family means first averaged endpoints within reporter and family; reporter–family values were then summarized. Intervals were estimated by reporter-level bootstrap resampling with fixed random seeds.

Reporter value was defined from the source study’s perturbation-distinctiveness analysis. For each mapped targeted marker, we averaged copairs mean average precision (mAP) across 1,001 perturbation labels (1,000 gene knockouts and NTC). Guide-level profiles generated by different sgRNAs for the same perturbation were positives and profiles from other perturbations were negatives, using cosine distance. To quantify undirected phase–marker overlap, we formed a 53 *×* 1,001 matrix containing phase and the 52 mapped marker distinctiveness profiles. For profile *r* and perturbation *g*, term frequency was TF_*rg*_ = *M*_*rg*_/ ∑_*h*_ *M*_*rh*_; document frequency was the number of profiles for which *M*_*rg*_ exceeded the across-profile mean for perturbation *g*; and IDF_*g*_ = log {(53 + 1)/(df_*g*_ + 1)} . Overlap was the Pearson correlation across perturbations between the TF–IDF-weighted phase and marker profiles. Directed recoverability was the unweighted median across ten methods of held-out-gene knockout-response Pearson. As a sampling-depth sensitivity analysis, we used one source-study subsampling draw that retained at most 300 cells per sgRNA without replacement, reaggregated guide-level profiles and recomputed perturbation-distinctiveness mAP. All 52 reporters mapped one-to-one to both source tables. Associations were Spearman correlations across reporters; 95% intervals used 3,000 reporter-level bootstrap resamples. Reporter value and cross-modal overlap were kept separate from recoverability because neither is a directed prediction metric.

### Same-cell pairing and within-condition controls

The primary same-cell analysis fixed the reporter-specific MLP and changed only the information available during fitting. The design covered 52 reporters, six intervention arms and five held-out-gene partitions, yielding 1,560 reporter–arm–partition results. Reporter was the inferential unit.

Exact pairing used the registered phase vector and targeted phenotype from the same cell. For gene–screen derangement, target vectors in the training and validation sets were reassigned without fixed points among cells from the same gene and screen; the held-out test set retained its true exact pairing. The model was therefore evaluated on the same exact-linked test outcome after its access to same-cell correspondence during fitting had been removed. For covariate-matched derangement, cells were matched without replacement within each gene–screen group by greedy nearest-neighbour pairing in standardized log cell area, log field–tile cell density and eccentricity. Density was the number of exact-linked cells in the same screen, well and imaging tile. These three outcome-independent phase covariates were scaled over the reporter cohort solely to construct the matching control; this transductive distance normalization used no targeted phenotype. Candidate pairs were ordered deterministically by distance and exchanged their target vectors. When an odd cell remained, it and the nearest existing pair were reassigned in a randomly oriented three-cycle; singleton groups were left unchanged. Training and validation partitions were deranged independently, whereas test cells retained exact pairing. Size–shape-only prediction restricted input to gross geometry features, whereas phase-minus-size–shape removed those features from the complete 172-dimensional representation.

The residual analysis addressed a different estimand. Within the training, validation and test partitions separately, a fixed-seed random permutation divided each gene–screen stratum into min(5, *n*) near-equal blocks. For cell *i* in block *b*, the target residual was

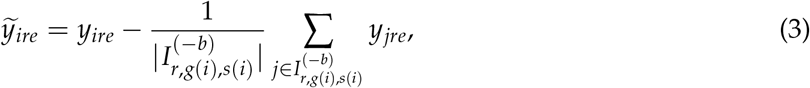

where the reference mean used cells in the same stratum but outside block *b*; singleton strata were assigned zero residual. The MLP was fitted directly to these residual targets. Because defining a test residual uses target measurements from peer cells in the held-out test stratum, this arm is a conditional association analysis rather than a deployable out-of-distribution prediction task; knockout-level recovery is consequently undefined.

Each reporter–arm score was averaged over the five held-out-gene partitions. Paired differences were calculated within reporter, and reporter-resampled intervals were used for mean exact-pair advantages. The residual arm was evaluated only at the cell level.

### Fidelity limits and sources of recoverability failure

For endpoint-wise recoverability, fold-specific training-set standardization does not affect Pearson correlation. Quantities that combine endpoint coordinates across held-out folds require a single reference scale. For reporter endpoint *e*, we therefore defined a common control scale from all unique non-targeting-control cells after removing the physical-screen-specific control mean,

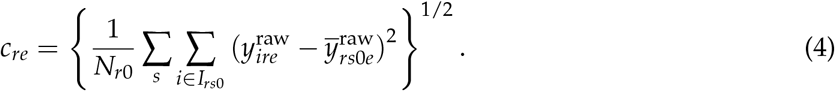

Here *N*_*r*0_ = ∑_*s*_| *I*_*rs*0_| . The common reference used control measurements only and was applied for evaluation, independently of model fitting and knockout outcomes. All 1,604 endpoint scales were finite and exceeded 10^*−*8^. If endpoint *e* had been standardized by training-fold standard deviation 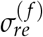, its control-relative response was converted as 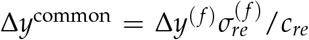, and predictions were converted identically. All ten methods shared the same fold-by-reporter target transforms. The original fold-standardized calculation was retained as a scale sensitivity analysis (Supplementary Methods). For each reporter and held-out gene, common-scale control-relative response vectors from different screens were then averaged with equal screen weight. Observed and predicted response magnitudes were

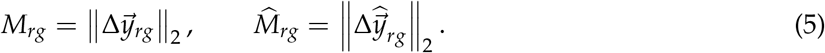

Magnitude-rank fidelity was corr_S_(*M*_*rg*_,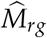 ), and the amplitude variance ratio was

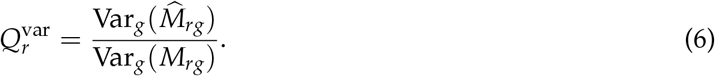

Strong-hit recall was the fraction of the 50 genes with largest observed magnitude recovered among the 50 genes with largest predicted magnitude. Profile fidelity was Pearson correlation between observed and predicted endpoint vectors within gene; it required at least three finite endpoint coordinates with non-zero variance and was otherwise undefined.

Reporter-level metrics were calculated from the element-wise unweighted median of ten aligned out-of-fold prediction profiles. The LysoTracker, ChromaLIVE 488 and pRb examples were selected to represent quantitative recovery, partial-pattern recovery and rank-preserving amplitude contraction, respectively, before final visualization.

### Target stability and coverage

Within-screen reliability used a deterministic hash of the phase-row identifier to split exact-linked cells into disjoint halves. Within each screen and half, reporter endpoints were robustly standardized to the non-targeting controls using the median and median absolute deviation. Endpoint responses were averaged by knockout and reduced to a root-mean-square response magnitude; Pearson correlation between halves was calculated over at least 20 common knockouts. Reporter reliability was the median correlation across its screens, and intervals used 1,000 gene-bootstrap resamples. Guide-half consistency used three predetermined partitions for knockouts represented by at least four guides. Endpoints were robustly standardized within well, aggregated with equal weight first across each guide subset and then across wells, and compared as knockout response magnitudes; a matched null used 100 random cell-half partitions within gene and well. Cross-screen consistency correlated complete-screen knockout-magnitude vectors for every pair of screens sharing a reporter and took the median pairwise correlation; it was estimable for 34 repeated reporters. Exact-link coverage was the number of linked reporter observations and was log_10_ transformed for modelling.

Bivariate associations used every reporter with the relevant measurement. A prespecified complete-case linear model included within-screen reliability, guide-half consistency and log_10_ exact-link coverage (*n* = 35). Cross-screen consistency was reported separately because it was available only for repeated reporters. The in-sample *R*^2^ was evaluated against 3,000 permutations of recoverability values across reporters while the three predictors and complete-case set were held fixed. Because the predictors overlap biologically and statistically, they were interpreted jointly as stability constraints rather than independent causal effects established by the associations.

For pRb and NPM1, response agreement was computed for three well pairs, three deterministic guide-half partitions and 100 random-cell-half partitions. Random-cell splits assess the sampling stability of aggregated responses within the observed population. Well and guide comparisons assess agreement across wells and perturbation reagents, respectively, combining sampling, biological and technical variation. The primary repeat comparison should match the intended use; alternative splits provide complementary evidence rather than isolate individual noise sources. Registered pRb microscopy used outcome-agnostic medoid cells from the same guide across three wells and served only to orient the experimental context.

### Environmental transfer

For each directed whole-screen evaluation, source controls comprised pooled cells from all other screens containing that reporter; source and destination groups were each deterministically limited to 20,000 cells. For *d* phase features or reporter endpoints, non-targeting-control shift was

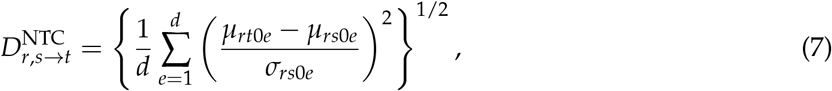

Coordinates with non-finite values or source-control standard deviation no greater than 10^*−*8^ were excluded, where *s* denotes the pooled source and *t* the destination screen. Transfer loss was the median-across-methods held-out-gene score minus the corresponding strict destination-screen score. Associations were quantified over 81 directions using Spearman correlation. For inference, phase shift, target shift and transfer loss were first summarized by their medians within each of the 34 reporters; two-sided *P* values were obtained by permuting the reporter-level shift summaries 3,000 times. Phase-control shift can be estimated before acquiring the destination targeted readout, whereas target-control shift is a retrospective explanatory quantity that requires that readout. A reporter was flagged as domain sensitive when its median cross-screen target stability was at or above the median among the 34 repeated reporters while its median strict-screen-minus-held-out-gene score was in the lowest quartile; the flag therefore marks an otherwise stable target with an unusually large model transfer penalty. Registered FeRhoNox control-cell images were deterministic phase and geometry medoids selected without reference to transfer performance.

### Cross-screen reporter stability

The context-stability analysis used all 81 frozen directed reporter-to-destination-screen evaluations for the 34 reporters observed in more than one screen. Recovery and magnitude-rank agreement were assessed separately. For each reporter and metric, all destination values at or above 0.70 were classified as stable above the gate, all values below 0.70 as stable below, and an observed range spanning 0.70 as crossing the gate. Supplementary Figure 4b shows the destination values, reporter median and observed range. This analysis did not recalculate the complete five-gate tier for each destination screen.

### Conditional ambiguity

Conditional ambiguity was evaluated for FeRhoNox in two screens and pRb in one screen using 80,975 exact-linked, non-control cells. Reporter–screen–gene groups with fewer than six cells were excluded. The 172 phase variables were standardized within group, and each cell was assigned its nearest other cell by root-mean-square Euclidean distance; where multiple guides were available, the neighbour was required to carry a different guide. Target endpoints were standardized to the non-targeting-control median and median absolute deviation within reporter, screen and well, then clipped to [*−* 10, 10]. Phenotype discrepancy was the root-mean-square end-point difference. For each gene, the median discrepancy of the directed morphology-nearest pairs was divided by the median discrepancy of all eligible unordered pairs. Values near one indicate that close morphology matching leaves most targeted-state discrepancy intact. A 32-dimensional principal-component representation of raw phase thumbnails, fitted separately within each reporter and screen, supplied a secondary comparison of the morphology-nearest pairs.

### Counterfactual response replacement and measurement-sufficiency classification

For each reporter, counterfactual response replacement substituted held-out knockout profiles with the element-wise unweighted median of the ten aligned predictors. The same downstream workflow was applied to measured and predicted profiles. The primary strong-hit set comprised the 50 largest response magnitudes among 1,000 knockout genes. Gene Ontology biological-process and cellular-component terms and Complex Portal complexes with 3–250 assayed members were tested by hypergeometric enrichment ^31–33^. Ranked-term overlap was the Jaccard index of the ten terms with smallest unadjusted *P* values, including nonsignificant terms.

We evaluated significant enrichment separately, using the 1,000 assayed genes as the background and Benjamini–Hochberg (BH) adjustment across all eligible terms, including zero-hit terms, within each reporter, annotation family, hit budget and measured or predicted list. Significant sets used adjusted *P ≤* 0.05. Significant recall was shared terms divided by measured significant terms and was undefined when the measured set was empty; significant Jaccard was undefined when both sets were empty. Ranking ties were resolved by full term name after *P* value. The cohort-relative downstream-retention index used an alternative ranked-list convention: 1,001 source-atlas labels, including NTC, with ties resolved by annotation source order. A sensitivity comparison found that changing the background alone left all 156 primary ranked-list comparisons unchanged, whereas changing tie order changed 20. Figure 5f,g and the significant-term analysis use the assayed-gene background and term-name ties. Fixed sensitivities used 5%, 10% and 20% hit budgets and lists of 5, 10 or 20 terms, preserving the same annotations and predictions (Supplementary Data 1).

Let *P*_52_(*z*_*r*_) denote the average-tie percentile rank of quantity *z* among the 52 reporters. The cohort-relative downstream-retention index was

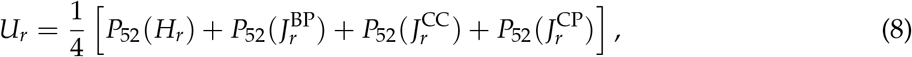

where *H* is top-5% hit recall and the three *J* terms are source-order top-ten overlaps for Gene Ontology biological process, Gene Ontology cellular component and Complex Portal. Magnitude ranking was excluded from *U*_*r*_ so that it could be evaluated as a separate predictor of downstream-analysis retention.

To relate fidelity to retained utility, we fitted Ridge regression using recoverability, magnitude-rank fidelity, magnitude-variance fidelity and within-screen reliability. Magnitude-variance fidelity transformed the variance ratio in Eq. 6 to 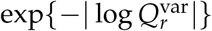, assigning the same value to reciprocal variance ratios. This transformation measures preservation of the spread of knockout-response magnitudes; it is not an intercept-and-slope calibration analysis. Evaluation used strict nested leave-one-reporter-out cross-validation. In each outer split, the four utility components for the 51 training reporters were converted to average-tie percentile ranks within that training set. Each held-out raw component was mapped to the corresponding training empirical distribution, with values outside the training range mapped to 0 or 1 and ties assigned the mean training percentile. The Ridge penalty was then selected in an inner leave-one-reporter-out loop that repeated the same training-only target construction. The held-out reporter’s downstream metrics were used only to construct its evaluation target against the training reference distributions; they did not determine those distributions or model selection. Predictive agreement was summarized by outer-loop Spearman correlation and *R*^2^; univariate reporter-level associations used the same strict outer targets.

The recovery–utility association was evaluated using Spearman correlation across reporters. For rank statistics and their bootstrap intervals in Supplementary Figure 4a, utility values were rounded to 12 decimal places before assigning average ranks, preserving mathematical percentile ties against floating-point perturbations; the underlying values and predictions were unchanged.

Candidate Ridge penalties in the nested analysis were *α* ∈{0.01, 0.1, 1, 10, 100} and were selected by mean inner-loop squared error.

### Measurement-sufficiency classification

An evaluation object was indexed by target, predictor, representation, cellular context, aggregation unit, task and tolerance. Response-fidelity tiers summarized reconstruction of a specified response profile; task decisions assessed a separately defined use. A quantitative proxy met the specified correlation, magnitude-spread and hit gates; this category did not estimate per-endpoint calibration slopes or intercepts. Not identifiable denoted insufficient reference evidence for the assessment, with low reliability and missing reliability recorded separately. Unresolved included missing fidelity evidence or complete evidence that satisfied neither proxy nor measurement-required criteria.

Rules were selected on this atlas, fixed before final tier assignment and applied hierarchically. These tiers concern response-profile fidelity and perturbation prioritization and were generated directly from the following fidelity and reliability criteria; the cohort-relative utility index in Eq. 8 was not used for tier assignment. A reporter was first classified as not identifiable when within-screen reliability was unavailable or *<* 0.30; otherwise, missing required fidelity evidence yielded unresolved. Among reporters with complete evidence, a quantitative proxy required held-out-gene recoverability *≥* 0.70, magnitude Spearman *≥* 0.70, variance ratio from 0.50 to 1.50 and top-5% recall *≥* 0.60. A ranking proxy required magnitude Spearman *≥* 0.70 and top-5% recall *≥* 0.50 but did not satisfy all quantitative criteria. Measurement required was assigned next when recoverability was *<* 0.60; all remaining reporters were unresolved. This precedence allows a reproducible target that preserves perturbation ranking to remain a ranking proxy even when its recoverability falls below 0.60. The cut-offs are descriptive operating points for this atlas and intended uses, not cross-system standards; their local sensitivity is reported in Supplementary Table 5. For a new application, tolerances should be specified from the intended decision, error costs and relevant repeat measurements using development data, then evaluated on held-out perturbations. Domain sensitivity was recorded as a separate whole-screen-transfer flag and did not constitute a sixth tier.

To assess predictor dependence, we recalculated all four response criteria from each method’s own predictions, using the same checkpoint states as the response-replacement ensemble, and reapplied unchanged reliability, thresholds and precedence. We distinguished quantitative–ranking transitions from changes across the proxy-use boundary (quantitative or ranking versus measurement required or unresolved). The primary denominator was the 43 reporters with reliability at least 0.30; the nine reliability-gated assignments were reported separately. Single-model benchmark states supplied a second complete comparison. Complete evidence and prediction-state sensitivity are provided in Supplementary Data 1 and Supplementary Methods.

### Calibration–evaluation assessment of measurement-use decisions

For a fixed predictor, calibration perturbations supplied acceptance evidence and disjoint evaluation perturbations supplied the task outcome. If *A* indicates acceptance and *S* evaluation success, the four outcomes were supported success (*A* = 1, *S* = 1), supported failure (1, 0), missed success (0, 1) and true rejection (0, 0). We reported coverage (*N*_11_ + *N*_10_)/*N*, false-support fraction *N*_10_/(*N*_11_ + *N*_10_), and success recall *N*_11_/(*N*_11_ + *N*_01_). A zero denominator was undefined. Missing calibration metrics could not satisfy a gate; missing evaluation outcomes were excluded from performance denominators and reported separately. All primary cases in the three decision ledgers had estimable evaluation outcomes. Supplementary Tables 8 and 9 specify the complete rules, study units, budgets and random-selection references.

### OPS decision calibration

We reused frozen outer-fold predictions, with a reporter-specific MLP as the primary predictor and Ridge, scButterfly and the fixed ensemble as sensitivities. The original model used outer fold *k* for testing, (*k* + 1) mod 5 for validation and the other three folds for fitting. Within each outer test fold, complete genes were partitioned into calibration and evaluation halves using seed 20261008. Calibration/evaluation gene counts were 105/106, 93/94, 91/92, 98/99 and 111/111; every guide and screen observation of a gene stayed together. Both halves were unseen by that fold’s fitted predictor and model-selection procedure. Profiles were converted to the common NTC endpoint scale (Eq. 4) and averaged equally across screens before response magnitudes were computed. Predictions from different outer-fold models were not pooled within a calibration case.

The primary task selected the largest absolute responses at a 20% budget, rounded up, and required evaluation hit recall *≥*0.60. Recovery-only accepted calibration *r≥* 0.70; rank-only accepted magnitude Spearman *ρ≥* 0.70; rank-plus-hit required both *ρ ≥*0.70 and calibration task recall *≥*0.50. Full-response required *r≥* 0.70, *ρ≥* 0.70, variance ratio 0.50–1.50 and calibration task recall 0.60. Reference reliability was not a gate in this all-52-reporter decision experiment; the 43 reporters passing the atlas reliability criterion supplied a separate sensitivity cohort. An additional full-response rule used the atlas tier’s 5% calibration hit gate while evaluating the 20% task.

Supported-use gain for rules *a* and *b* was 100(*N*_11,*a*_*− N*_11,*b*_)/*N*, in percentage points of all cases. Twenty seeds (20261008–20261027) changed only the calibration/evaluation allocation, not model fitting or gate thresholds. Fixed secondary analyses changed selection budgets to 5% or 10% and success thresholds to 0.50, 0.70 or 0.80. For matched coverage, calibration recovery, rank, task recall or a minimum normalized gate margin ranked reporters within each predictor–fold–seed. With calibration recovery *r*, magnitude rank *ρ*, variance ratio *v* and hit recall *h*_*b*_ at selection budget *b*, the combined-rule scores were

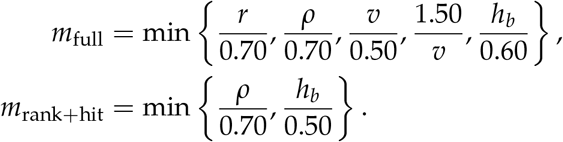

Each component is a ratio to its gate boundary, oriented so that values at least one satisfy the gate; the minimum records the least-satisfied condition. Reporters were ranked by descending score, and the highest *cn* were accepted for *c* = 0.25, 0.50, 0.75, 1.00, with reporter identifier breaking ties. The primary analysis comprised 52 reporters times five folds. Intervals used 2,000 paired reporter-cluster bootstrap draws, keeping every fold and partition of a sampled reporter together. These intervals condition on the observed perturbations and frozen predictors.

### Independent applications

We surveyed public imaging and perturbation repositories and associated study deposits, recording 59 candidate resources by 12 August 2026. Each entry recorded six capabilities—a lower-cost broad phenotypic input, a targeted readout, paired observations, perturbations, within-screen negative controls and repeated acquisition of the targeted readout—together with the first capability that determined eligibility and the strength of the supporting evidence. Forty-two resources could be classified from the available evidence. Two resources (OASIS and PERISCOPE) met the full design criteria for this audit; Cell Health and LINCS supplied separate, task-specific prediction and decision ledgers analysed with the same fixed operating points. Aggregate audit outcomes are in Supplementary Table 1; the complete 59-row registry with evidence and exclusion reasons accompanies the source data.

The first selected resource, cpg0037-oasis, comprised 66 plates and 21,695 wells from a commercial five-donor pool of primary human hepatocytes exposed to 1,085 non-control compounds ^35^. Pairing was at the well rather than same-cell level. Inputs were 808 CellProfiler features attributable to the brightfield channel; 120 brightfield–fluorescence cross-channel features, 4,512 fluorescence-attributable features and 200 channel-free features derived from segmentation were excluded. The retained features used fluorescence-derived object masks, making prediction conditional on the deposited segmentation. Primary targets were the deposited plate-normalized metabolic-activity and lactate-dehydrogenase-release endpoints. Compounds were randomly assigned to five outer folds with all wells and concentrations of a compound held out together. Outer-training-set median imputation and standardization preceded Ridge regression with fixed *α* = 1. Recovery used the deposited plate-normalized endpoints. For response-fidelity analysis, observed and predicted well values were averaged by compound, plate and reporter, centred on the corresponding within-fold, plate-specific negative-control mean and then averaged across plates by compound. Reliability was Pearson correlation across compounds between two seeded, disjoint half-means of replicate non-control wells pooled across plates and concentrations, yielding a dose-averaged reference.

### Cell Health

We used fluorescent Cell Painting profiles and separately acquired cell-health endpoints from ES2 and HCC44^15^. Pairing was by cell context and guide, not by cell or assay well. Each context contained 49 eligible genes and 100 targeting guides with identifiable mappings. Endpoint-specific inclusion required finite targets on both reference plates: 100 guides met this condition for nine endpoints and 94 for ROS. Ambiguous mappings were quarantined; the A549 context was excluded because duplicate keys could not be resolved. Ten endpoints were specified before fitting: G1, S, G2 and mitotic fractions; dead, early-apoptotic and late-apoptotic fractions; live-cell count; ROS; and H2AX. These produced 20 endpoint–context cases.

Within each assay, wells were averaged by guide and plate, centred on the median of ten infected non-targeting guides, and then averaged equally across plates. Empty-well controls were excluded. Cell Painting predictors contained 1,783 numeric features and no identifiers. Imputation, constant-feature filtering and standardization used training genes only; targets were never imputed. Training-gene weighted means and standard deviations standardized each target, and predictions were converted back to control-relative units. The 49 genes were shuffled with seed 20261008 into 25 training, 12 calibration and 12 evaluation genes, grouping all guides and both cell contexts by gene. A separate Ridge predictor with fixed *α* = 100 was fitted for each endpoint–context pair, using equal total weight per gene. Model parameters were not selected on decision-calibration genes.

The task used the top 30% absolute control-relative responses, rounded up to four genes in each held-out half. Both task acceptance and evaluation success required recall *≥* 0.60, or at least three of four hits. Recovery-only required calibration signed Pearson *r≥* 0.70; rank-only required absolute-response Spearman *ρ ≥* 0.70; task-plus-repeat added calibration Pearson agreement *≥* 0.30 between the two target plates. A secondary repeat-supported outcome required success separately against each plate, rather than only their mean. Twenty fixed gene partitions (seeds 20261008–20261027) assessed allocation sensitivity. The endpoints share cells, pathways and reference assays; partition repetitions were not independent biological replicates. Figure 6c displays the original fraction changes on symmetric-log axes (linear within *±*0.01); metrics and hit selection use the untransformed responses.

### LINCS

The matched LINCS-Pilot1 resource links A549 Cell Painting at 48 h with separately acquired L1000 RNA at 24 h ^22^. We retained actual 10 *µ*M conditions with at least two image plates and two RNA plates, excluding DMSO, MG132 and bortezomib from prediction targets. The eligible cohort comprised 1,292 compounds in 1,286 chemical-identity groups and 978 landmark RNA probes (957 uniquely annotated and 21 ambiguously annotated). Chemical identities were grouped by the first 14 InChIKey characters, falling back to compound identifiers. The primary partition assigned 775 compounds to training, 257 to calibration and 260 to evaluation, grouping all doses and replicate observations by chemical identity.

Image features used plate-DMSO median/MAD normalization and RNA values used plate-DMSO centring. Wells were averaged within compound–plate and then equally across plates in each assay, without pairing all image wells with all RNA wells. RNA references A and B used odd and even X-acquisition groups, respectively; the consensus averaged all eligible plates and therefore included both references. The input schema retained 1,663 shared numeric morphology features after excluding coordinates, identifiers and nonshared columns. Training compounds determined median imputation, 1st/99th-percentile clipping, constant-feature removal and input/target standardization.

Ridge selected *α* ∈ {10, 100, 1000, 10000} by three-fold chemical-group cross-validation within training. The MLP used 256/128 ReLU units, dropout 0.1 and AdamW (learning rate 0.001, weight decay 0.01, batch size 128), with at most 200 epochs and early-stopping patience 25. A training-only 20% validation split selected the epoch before refitting on all training compounds. Six rules were evaluated: accept all; recovery (*r≥* 0.70); magnitude rank (*ρ ≥*0.70); task (top-10% absolute-response recall*≥* 0.60); task plus repeat (calibration A/B signed Pearson *≥*0.30); and task plus repeat plus rank. Evaluation success required at least 16 of 26 measured hits. Repeat-supported success additionally required this overlap with both A and B. Ten fixed chemical-group partitions assessed Ridge sensitivity. Two hundred joint evaluation-chemical-group bootstrap draws preserved endpoint dependence and recomputed hit sets with calibration decisions fixed. No endpoint-specific threshold was selected from evaluation results.

### PERISCOPE

The cpg0021-periscope resource paired DAPI, ConA, phalloidin and WGA Cell Painting measurements with anti-TOMM20 fluorescence in nine plates of A549-TetR-Cas9 cells ^10^. It was analysed at two resolutions: 1,080,000 single cells obtained by a deterministic cap of the first 120,000 streamed rows per plate, and all 704,771 deposited median-aggregated gene–sgRNA–plate profiles. Inputs comprised 2,508 measurements attributable exclusively to the four fluorescent predictor channels and targets comprised 823 anti-TOMM20-associated measurements; 60 predictor–target cross-channel and 375 channel-free features were excluded. Segmentation used phenotypic images that included the target channel, making this a cross-channel prediction analysis conditional on the deposited masks. Non-control genes were assigned to five outer folds; the next cyclic fold provided inner validation. Nontargeting controls remained in training. The Ridge penalty was selected by endpoint-standardized validation MSE and the model was refitted on the complete outer training set. Within each test fold, observed and predicted measurements were averaged by gene and plate and centred on the same-plate nontargeting-control mean. Recoverability was then calculated by dividing each endpoint’s observed and predicted responses by its observed response standard deviation and pooling the gene–plate–endpoint coordinates for Pearson correlation. This endpoint-standardized pooled statistic differs from the equal-endpoint macro-average used for OPS (Eq. 2); endpoint inclusion and response-magnitude scaling are detailed in Supplementary Methods. Reliability split the sgRNAs for each eligible gene into two disjoint halves within plate, centred each half on plate-specific controls, standardized endpoints by the full-guide response spread and summarized the nine plate-level correlations by their median; genes with fewer than two sgRNAs and negligible-spread endpoints were excluded. This estimate concerns reproducibility of guide-defined gene responses and combines measurement, guide and biological variation.

The atlas response-fidelity tier rules were applied to the OASIS CellProfiler and PERISCOPE assessments without re-estimation. We assessed the dependence of these classifications on the numerical cut-offs using the same 36 threshold variations as in OPS, with predictions and metric aggregation unchanged (Supplementary Table 7 and Supplementary Data 2). Batch-specific analyses, reliability sensitivity, leave-one-dye-out analyses and within-screen input-permutation controls are described in Supplementary Methods.

#### Brightfield-guided metabolic-activity prioritization

The focused OASIS analysis used 768 features from the released brightfield-channel DINO representation and the deposited plate-normalized RealTime-Glo metabolic-activity readout. A fixed compound partition comprised 868 development compounds (13,157 wells) and 217 test compounds (3,300 wells). All 868 development compounds contributed to fitting; Hycanthone had seven rather than eight assayed concentrations and was excluded only from development out-of-fold compound scoring, leaving 867 compounds for that comparison. All test compounds had eight assayed concentrations. All measurements of test compounds were excluded from fitting across doses and production sources. The partition was inherited from a preceding representation exploration. The test compounds had not been evaluated in that brightfield-DINO exploration, although their target measurements were part of the public resource and its earlier CellProfiler analysis. The target, predictor settings, signed scoring rule and selection budgets were fixed before final fitting and held-out scoring. Brightfield inputs were centred on same-plate DMSO means, followed by training-only median imputation and standardization. Ridge regression used *α* = 100 and all 768 features, without compound identity, concentration, fluorescence or cell-count inputs. Observed and predicted responses were each centred on their corresponding same-plate DMSO mean.

We defined loss as the negative control-relative response, so an increase in metabolic activity did not count as a loss. Within each compound and actual concentration, wells were averaged within production source, then equally across available sources; these values were averaged equally across the eight assayed concentration positions. This score concerns the measured concentration panel, whose range can differ between compounds, rather than a common-range dose–response area. The primary selection budget was the top 5%, rounded up to 11 of 217 compounds; fixed secondary budgets were 10% and 20%. The 94-compound complete-source subset supplied dose-matched, cross-source prediction and measured-repeat comparisons, with five compounds selected at the primary budget. Compound bootstrap intervals used 2,000 resamples that preserved the dose and source pairing. They condition on the fitted predictor and observed sources. Predictor sensitivities, correspondence controls and complete source coverage are described in Supplementary Methods; predictions and selection records accompany Supplementary Data 3. The released representations came from bright-field channels of stained, fixed-cell acquisitions.

#### Label-efficiency and raw-phase representation analyses

##### Label-efficiency analysis

The low-label analysis (Supplementary Figure 2a) used a panel fixed before the learning-curve analysis—pS6, NPM1, SEC23A, LAMP1, 5xUPRE, TOMM20, FastAct, EEA1, WGA, Peroxi, Hoechst and G3BP1—and treated the remaining 40 reporters as a fully labelled donor atlas. The panel spans the ten biological-system annotations and includes reporters across the full-label recoverability range; it was not selected by low-label performance. Exact target-reporter observations were sampled independently within control-status-by-gene strata by Hamilton largest-remainder allocation, yielding exact budgets of 0.1%, 1% and 20% without consulting target values. The same fraction was applied to training and inner-validation cells; outer-test cells remained complete and sealed, and common sampling manifests were used across the five held-out-gene partitions. Ridge, gradient-boosted trees, CatBoost, the reporter-specific MLP, TabM, scButterfly and scPair used only the sparsely labelled target reporters. The shared residual MLP, MultiTab and MIDAS additionally used complete donor-atlas supervision. These two groups therefore represent target-only and atlas-assisted deployment settings rather than an equal-information ranking of algorithms. Equal-reporter partition means and standard deviations were calculated at each fraction. Matched 100% evaluations came from a separately completed full-label experiment on the same reporter panel and partitions. The dashed connection from 20% to 100% therefore shows a cross-experiment comparison. Reporter identities and selected row indices for every partition and fraction are provided in the released sampling manifests.

##### Raw-phase representation analysis

The raw-phase representation analysis (Supplementary Figure 2b) used six reporters—EEA1, LAMP1, TOMM20, NPM1, pS6 and G3BP1—and five held-out-gene partitions. The reporters were fixed before representation comparison to span six biological systems and the full-label recoverability range, subject to availability of registered raw crops; they were not chosen by image-model performance. Registered 384 *×* 384 level-0 phase crops were encoded with a fixed pretrained DINOv2 ViT-S/14 vision transformer ^44^, a fixed pretrained Cytoland VSCyto2D encoder, a partially fine-tuned Cytoland encoder or the engineered 172-dimensional representation ^45^. Cytoland used the released VSCyto2D checkpoint. Valid pixels in each crop were centred by their median and scaled by their interquartile range; padded pixels were set to the valid-pixel median and no image augmentation was applied. For partial fine-tuning, the stem and first two encoder stages were fixed, the final two stages and a 256–128 reporter endpoint head were trained for at most 25 epochs using AdamW, batch size 48 and validation-based early stopping with patience 6. Backbone and head learning rates were 10^*−*5^ and 3 *×* 10^*−*4^, respectively, with two warm-up epochs followed by cosine decay. Predictions were paired within each reporter and outer partition (6 *×* 5 = 30 reporter–partition results per representation). Matrix entries show five-partition means and standard deviations within reporter. Comparative inference used the six reporters, with paired mean differences and the deposited reporter-hierarchical 95% bootstrap intervals. The complete reporter and crop manifests accompany the source data.

##### Statistical analysis and reproducibility

Pearson correlation measured linear association between observed and predicted phenotypes or responses; Spearman correlation measured rank agreement. Confidence intervals were obtained by resampling the highest-level unit appropriate to the claim, generally reporter. Unless stated otherwise, bootstrap intervals were percentile 95% intervals from 10,000 deterministic reporter resamples. Reporter-to-screen analyses used reporter-level summaries or reporter-preserving resampling as specified above. Holm correction was applied to the prespecified reporter-level method-comparison family; BH correction was applied to enrichment terms within the families defined above. Individual association permutations and bootstrap intervals were not multiplicity-adjusted. Genes, endpoints, cells and partitions nested within reporter were not counted as independent reporter replicates. Paired Wilcoxon tests and correlation permutation tests were two-sided; the joint-model *R*^2^ permutation test used the upper tail of its null distribution. Exact sample sizes, exclusions, bootstrap draws and permutation counts are given in the figure legends and panel-level source tables. Random seeds and resampling draws are reproduced by the released workflows.

##### Software and hardware

Analyses used Python 3.10.20, NumPy 1.26.4, pandas 2.1.4, SciPy 1.11.4 and scikit-learn 1.7.2; neural models used PyTorch 2.3.1, gradient-boosted trees used XGBoost 2.1.4 and CatBoost used version 1.2.10. Computations used NVIDIA H100 GPUs (80 GB) and multi-core CPUs. Independent reporter- and fold-level jobs were run in parallel, with up to four concurrent jobs per GPU in the same-cell analyses. Figures were rendered with Matplotlib 3.7.5.

## Author contributions

Mengran Li, Jianqing Zhu and Bo Li conceived the study with Ronghui Zhang, Lian Zhang and Jinchao Xu. Mengran Li led method development, software, formal analysis, visualization and data curation. Jianqing Zhu and Bo Li contributed to methodology, validation, interpretation and project coordination. Chengyang Zhang, Zhenchao Tang, Jiaying Wang, Wenbin Xing, Boyu Zhang, Jin-feng Xu, Lingbei Meng, Bob Zhang and Junzhou Chen contributed to validation, interpretation and manuscript revision. Ronghui Zhang, Lian Zhang and Jinchao Xu supervised the work and contributed to project administration. Mengran Li, Jianqing Zhu and Bo Li wrote the original draft. All authors reviewed and edited the manuscript and approved the final version.

## Data availability

The optical perturbation atlas was generated by the OPS study ^27^. We used revision 6 of its public Zenodo record (https://doi.org/10.5281/zenodo.20495192), released 1 June 2026 under CC BY 4.0^46^. Full imaging and molecular data remain available through the Biohub OPS Explorer (https://biohub.ai/ops-explorer) under the source resource’s terms. The external resources cpg0037-oasis, cpg0021-periscope, cpg0004-lincs (Cell Painting) and cpg0003-rosetta (LINCS-Pilot1 L1000) are available through the Cell Painting Gallery ^10,22,35,47^. Cell Health inputs and targets were obtained from the public version 2.0 deposit (https://github.com/broadinstitute/cell-health/tree/v2.0) ^15^. Source-file hashes, LINCS RNA S3 version identifiers, image-feature manifests and chemical/guide matching records accompany Supplementary Data 3.

OASIS DINO profiles and metadata for metabolic-activity prioritization were obtained from the source study’s Axiom OASIS deposit (https://doi.org/10.5281/zenodo.17067683, Jessica Ewald, CC BY 4.0). Derived tables retain source attribution and the deposited numerical values; predictions, aggregation and selection records are outputs of the present analysis.

Processed OPS phase features, exact-linked reporter targets, metadata and fixed field- and gene-holdout assignments are available in the MorphoSuff dataset repository (https://huggingface.co/datasets/Amanda1998/MorphoSuff). Supplementary Data 1–3 accompany this manuscript. They contain OPS predictor-specific evidence, calibration–evaluation decisions and functional analyses; external threshold sensitivity; and held-out OASIS, Cell Health and LINCS predictions and selection records, respectively. Full raw microscopy remains with the source resources; provenance and source-specific licences accompany the data.

Gene Ontology ^31,32^, Complex Portal ^33^, CEGv2^48^ and MitoCarta 3.0^49^ were reused as public third-party annotations. The releases, download dates and identifiers used in the analysis are recorded in the source-data provenance.

## Code availability

MorphoSuff software for data preparation, model fitting and measurement-sufficiency assessment is available at https://github.com/limengran98/MorphoSuff under the MIT License. Reconstruction and verification code for the reported decision, functional and external-use analyses accompanies Supplementary Data 1–3. The Python package provides dataset adapters and shared response-fidelity, reliability and measurement-decision analyses for use with other paired datasets or user-supplied predictions.

