## Supplementary Information for "When label-free morphology is sufficient for targeted cellular measurements"

#### Contents

- Supplementary Figures 1 to 6
- Supplementary Tables 1 to 9
- Supplementary Data 1 to 3
- Supplementary Methods
- Supplementary Notes 1 and 2
- Supplementary References

### Supplementary Figures

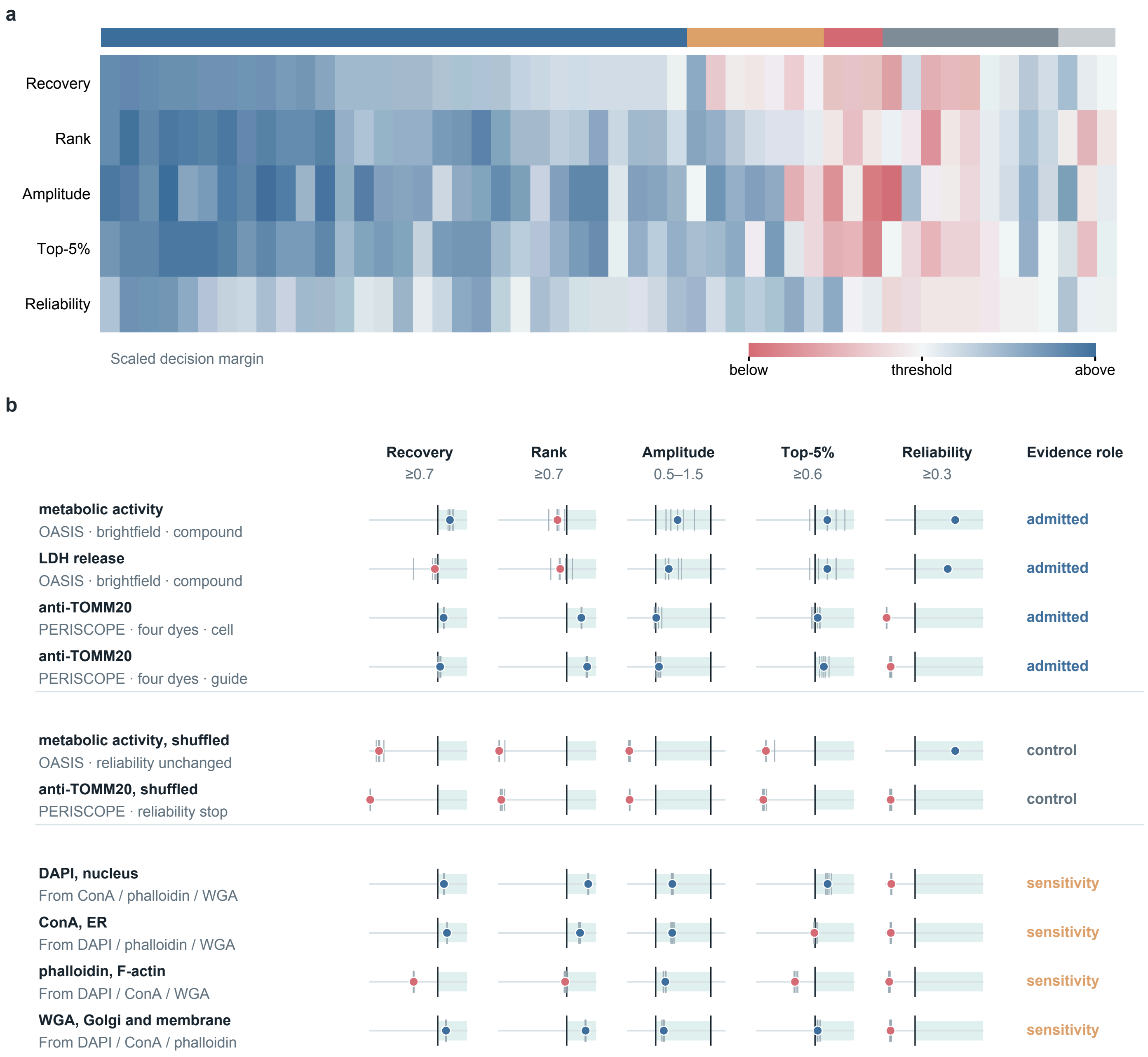


Supplementary Figure 1: **Measurement-sufficiency evidence across three experimental contexts. a,** All 52 OPS reporters evaluated against the five criteria. Columns are reporters ordered by decision tier; the upper strip denotes quantitative proxy, ranking proxy, measurement required, not identifiable and unresolved assignments (blue, orange, rose, grey and pale grey, respectively). Colour gives the signed, scaled margin to the corresponding criterion, with blue indicating that the criterion is met and red that it is not. Exact values, reporter identities and ordering accompany the source data. **b,** Evidence for the two independent measurement settings, their input-permutation controls and the four-dye substitution sensitivity analysis. Points show fold, screen or dye estimates; large markers are the reported summaries and vertical lines indicate thresholds. Dye substitutions address prediction between fluorescent channels, separately from the targeted-readout applications. Resource selection and OASIS batch-specific classifications are reported in Supplementary Tables 1 and 3, respectively.

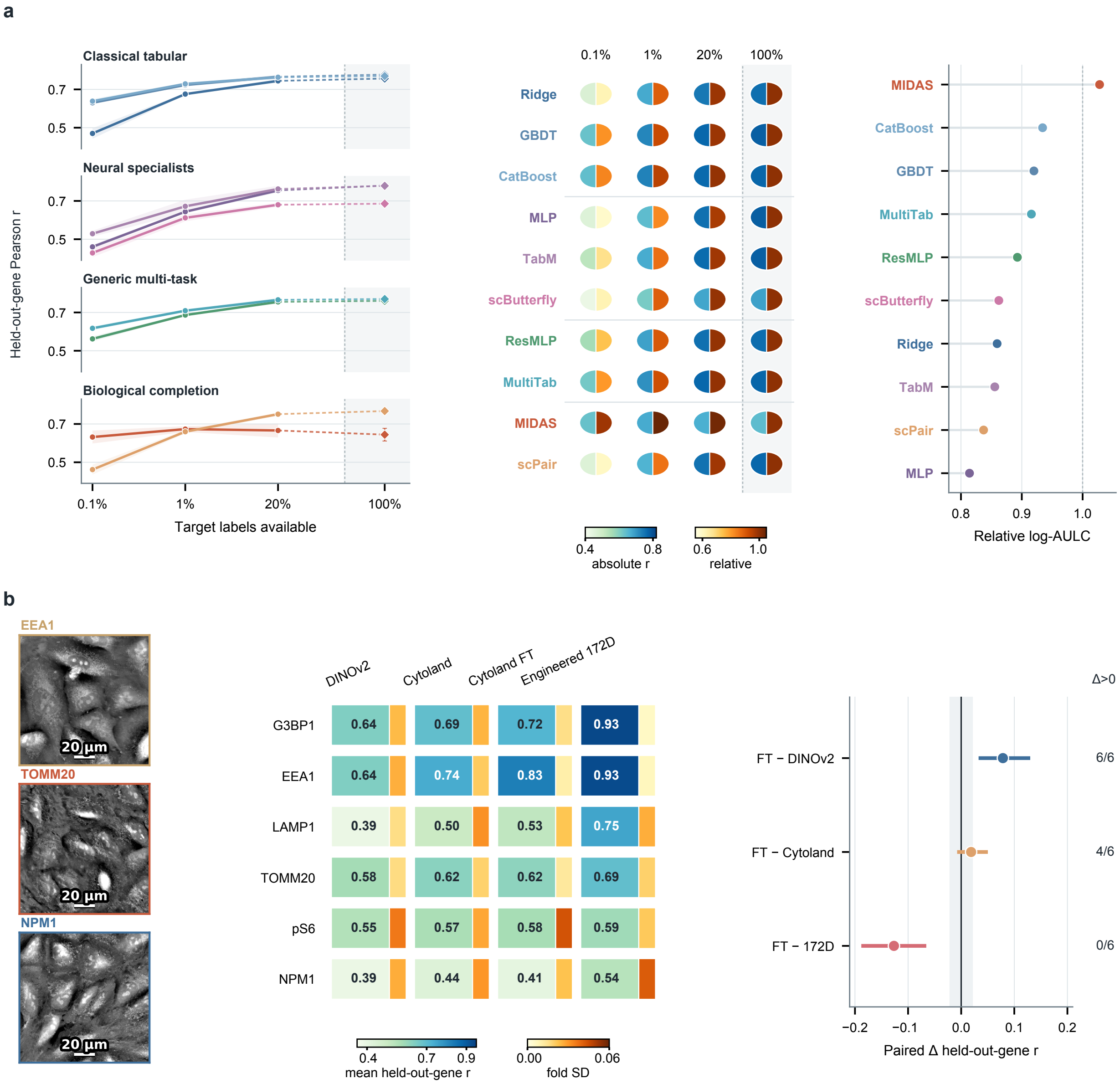


Supplementary Figure 2: **Label-efficiency calibration and representation sensitivity.** **a,** Held-out-gene performance on a panel of 12 reporters fixed before the learning-curve analysis. Curves and bands show equal-reporter means and one standard deviation across five outer gene partitions for ten methods at 0.1%, 1% and 20% target labels; diamonds are matched 100% evaluations from a separate full-label experiment, and dashed 20–100% segments connect the experiments. In each split glyph, the left half gives Pearson correlation and the right half gives recovery relative to the method's 100% result. Relative log-AULC is the trapezoidal mean of relative recovery over $\log_{10}$ label fraction from 0.1% to 20%. **b,** Raw-phase representation comparison for six reporters fixed before representation analysis. The matrix gives the five-fold mean held-out-gene Pearson correlation for each reporter and representation, and the adjacent strip gives the fold standard deviation. Paired reporter-level contrasts compare partially fine-tuned Cytoland with frozen DINOv2, frozen Cytoland and engineered 172-dimensional morphology; points are mean differences, intervals are reporter-hierarchical 95% bootstrap confidence intervals and counts denote reporters with a positive mean difference. Registered phase crops illustrate three inputs; scale bars, 20 $\mu$m. The full-label engineered-feature analysis supplies the reporter-level tiers.

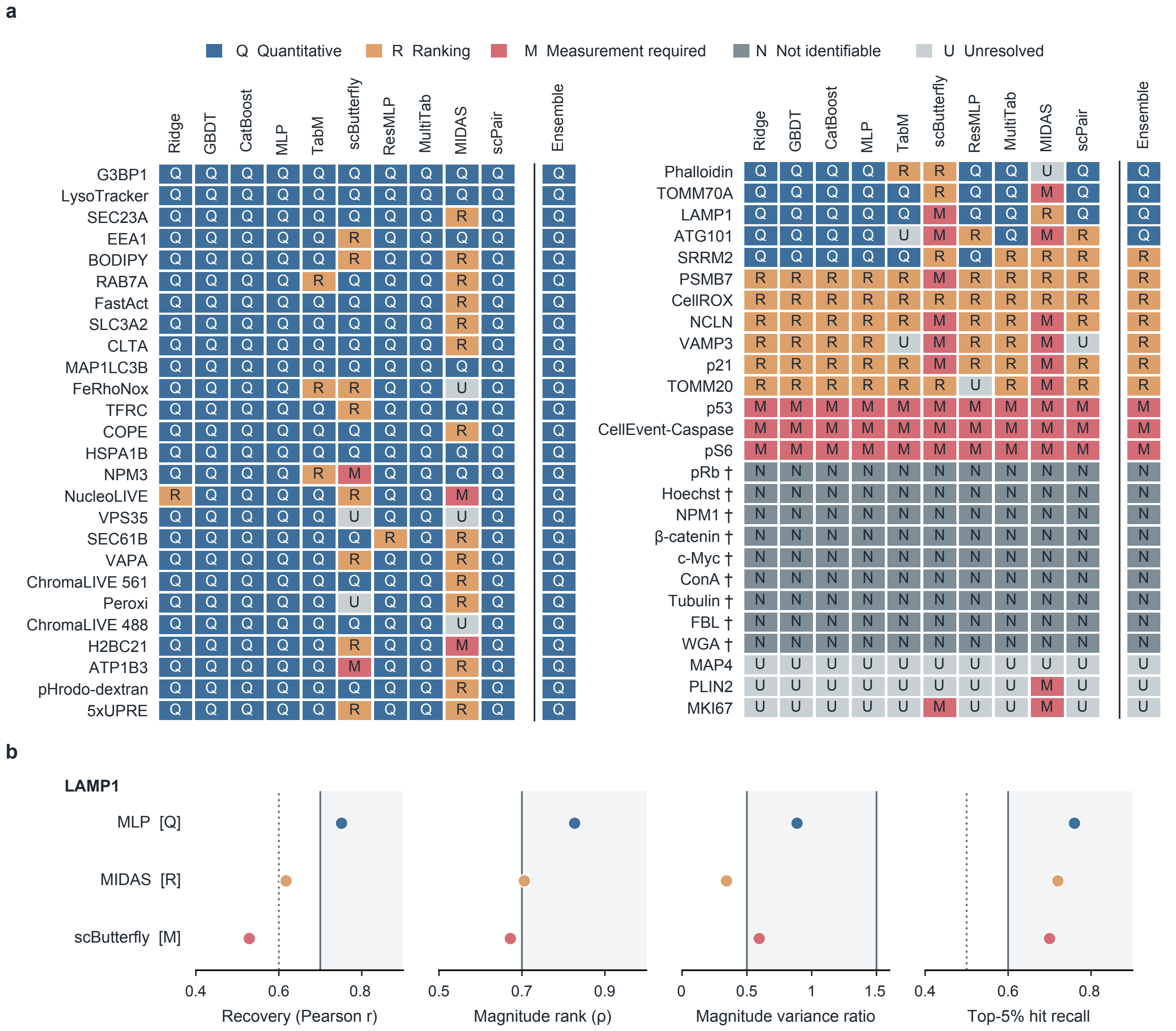


Supplementary Figure 3: **Predictor choice changes the measurement uses supported for individual targets. a,** Complete assignments for 52 reporters and ten predictors, with the fixed median ensemble as reference. The two blocks continue the reporter order used in Supplementary Figure 1a; colours and letters distinguish quantitative proxy (Q), ranking proxy (R), measurement required (M), not identifiable (N) and unresolved (U). Each method supplies its own recoverability, magnitude rank, variance ratio and strong-hit recall, using the same prediction states as the ensemble; partitions, common control scale, target reliability and thresholds are held fixed. Of 43 reliability-passing reporters, 20 support proxy use under all ten methods, 6 under none and 17 cross the proxy-use boundary. Daggers identify the nine reliability-gated reporters. **b,** LAMP1 illustrates the continuous evidence under MLP, MIDAS and scButterfly; reliability is 0.448 for all three. Points are predictor-specific metrics; pale bands and solid boundaries mark quantitative criteria, and dotted boundaries mark recovery 0.60 for the measurement-required rule and hit recall 0.50 for the ranking rule. Both prediction states, all reporter–method metrics and criterion-level changes accompany Supplementary Data 1.

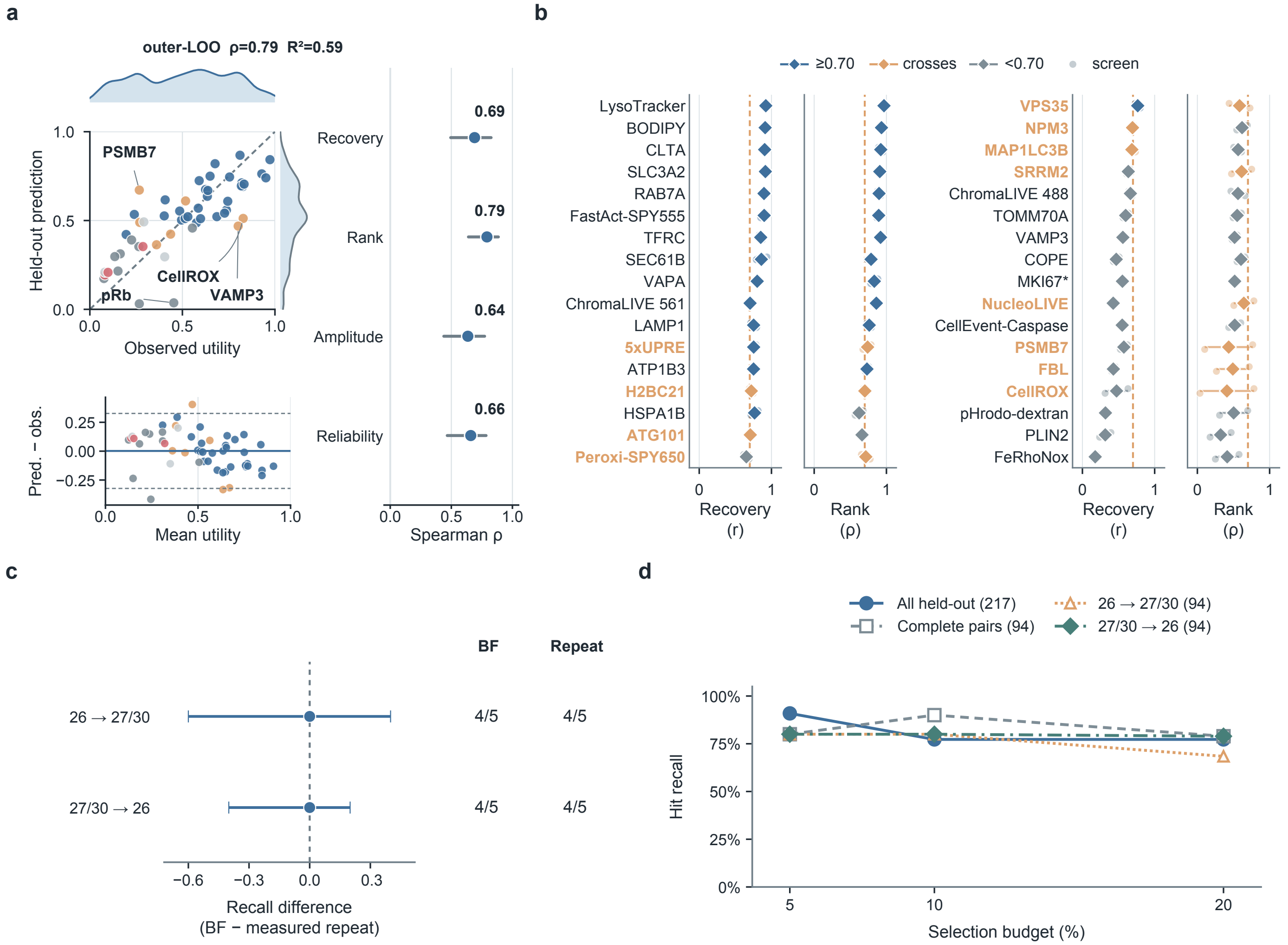


Supplementary Figure 4: **Utility prediction, screen context and repeat measurements refine use-specific assessment. a,** Nested leave-one-reporter-out utility prediction for 52 reporters, with training-only percentile references and model selection; Amplitude denotes magnitude-variance fidelity and forest intervals are 95% reporter-bootstrap intervals from 10,000 resamples. Residuals are predicted minus observed utility, with lines at mean bias and mean ±1.96 standard deviations. **b,** Benchmark ten-method score medians for recovery and magnitude rank across 81 directions nested in 34 reporters: points are destination evaluations, diamonds reporter medians and segments ranges, ordered down two consecutive blocks of 17 reporters. Blue remains above 0.70, grey below and gold crosses; complete destination-specific tiers were not recalculated. **c,** Among 94 hepatocyte-screen compounds with complete matched concentrations in two sources, both cross-source directions recovered four of five strongest measured metabolic-activity losses, as did measured-repeat prioritization; paired recall-difference intervals are 95% compound-bootstrap intervals from 2,000 resamples. Source roles 0 and 1 correspond to production source 26 and its paired source (27 or 30), respectively; these production sources were represented in training. **d,** Recall at fixed 5%, 10% and 20% budgets, retaining the primary brightfield predictor and four evaluation populations or directions. Complete external outcomes, source-pair comparisons and selection records accompany Supplementary Data 3.

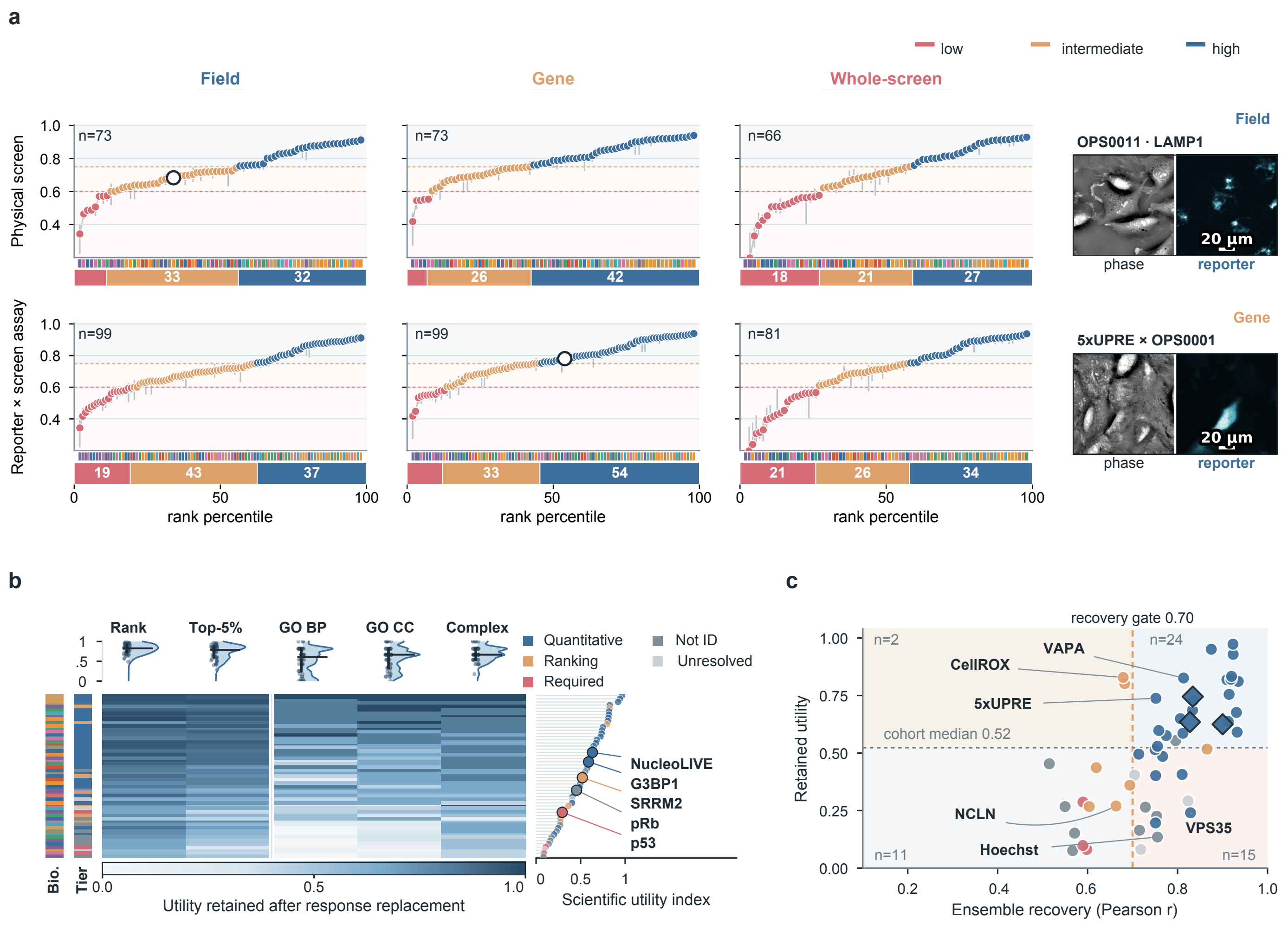


Supplementary Figure 5: **Screen-level recoverability and retained scientific utility provide complementary views of the atlas. a,** Knockout-response recoverability for physical screens and reporter–screen assays under field, gene and whole-screen holdouts. Points are unweighted medians across ten methods, ordered separately in each view; intervals span the method interquartile range, and bars count low ($r < 0.60$), intermediate ($0.60 \leq r < 0.75$) and high ($r \geq 0.75$) descriptive strata. Field/gene views contain 73 screens and 99 assays; whole-screen views contain 66 destination screens and 81 directed assays. Colour strips denote biological systems, and rings identify the outcome-independent exact-cell examples at right; scale bars, 20 $\mu$m. **b,** Retention of response ranking, strongest knockout hits and the ten highest-ranked functional terms after ensemble response replacement for 52 reporters, ordered by the utility index. Points, medians and interquartile ranges accompany the distributions; the utility index combines cohort-percentile hit and functional-list retention, and colours denote response tiers. **c,** Ensemble recoverability versus utility expressed relative to the leave-one-reporter-out training reference; diamonds identify domain-sensitive reporters. The recovery line at 0.70 is an operational criterion, whereas the horizontal line is the descriptive cohort median; ranked functional-list overlap in **b,c** is distinct from the significant-term analysis in main Figure 5g.

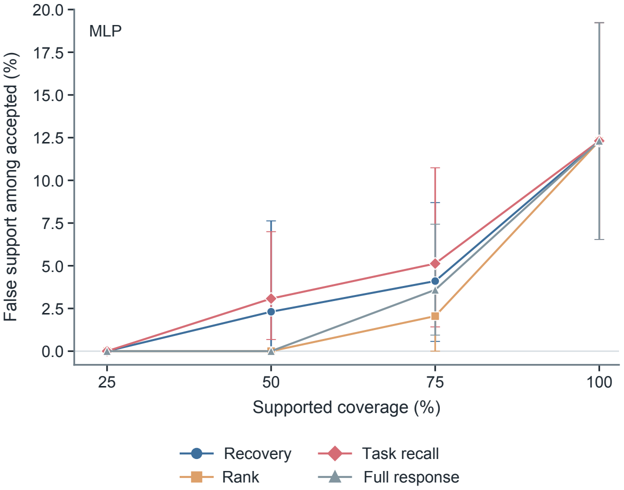


Supplementary Figure 6: **Task-support errors at matched coverage in held-out OPS perturbations.** Calibration recovery, response-magnitude rank, top-hit recall and full-response margin each selected the highest-scoring 25%, 50%, 75% or 100% of reporters within each of five outer folds. The same frozen MLP predictions, 52 reporters and primary calibration/evaluation gene allocation were used for all scores; plotted coverage reflects integer-rounded selection counts. Evaluation success required recovery of at least 60% of the strongest 20% of measured knockout responses. Points show the fraction of accepted reporter–fold cases failing that criterion, with 95% reporter-cluster bootstrap intervals from 2,000 resamples; each cluster retains its five folds. Lines connect the four predetermined coverage settings. All thresholds and selection ranks use calibration genes, with identifier-based resolution of score ties.

#### Supplementary Tables

Supplementary Table 1: **Assessment of public resources for measurement-sufficiency analysis.** Fifty-nine resources were considered and 42 could be classified from the available information. Counts give the first measurement-design or access condition determining each outcome. The two resources selected by the full eligibility audit support complementary applications: OASIS provides well-paired brightfield and biochemical measurements, and PERISCOPE provides paired fluorescent-channel measurements. Cell Health and LINCS are separate focused task ledgers and are not additional rows in this eligibility count. Resource-level eligibility records accompany the source data.

| Outcome | Measurement-design or access condition | *n* |
|---|---|---|
| Excluded | No lower-cost broad phenotypic input | 13 |
| Excluded | No targeted readout | 12 |
| Excluded | Incompatible pairing or no perturbations | 6 |
| Excluded | Restricted access, absent deposit or no cost advantage | 8 |
| Unresolved | Required measurements present; eligibility unresolved | 1 |
| Selected | OASIS and PERISCOPE | 2 |
| Not assessed | Information or accession records insufficient for classification | 17 |

Supplementary Table 2: **Measurement-sufficiency criteria applied to two independent public screens.** Thresholds were retained from the OPS analysis. Values are medians over five outer folds (held-out compounds for cpg0037 and held-out genes for cpg0021), except reliability, which is estimated separately from replicate measurements. Bold values fall outside the criteria for a quantitative proxy; classifications follow the hierarchy in Methods. Recoverability and reliability use the aggregation appropriate to each resource, as described in Supplementary Methods.

| Screen | Target (fitting unit) | $r$ | $\rho$ | Var. | Recall | Reliab. | Tier |
|---|---|---|---|---|---|---|---|
| cpg0037 | metabolic activity (well) | 0.826 | **0.604** | 0.897 | 0.727 | 0.716 | unresolved |
| cpg0037 | LDH release (well) | **0.670** | **0.633** | 0.736 | 0.727 | 0.638 | unresolved |
| cpg0021 | anti-TOMM20 (cell) | 0.760 | 0.852 | 0.509 | 0.627 | **0.005** | not identifiable |
| cpg0021 | anti-TOMM20 (guide) | 0.724 | 0.912 | 0.559 | 0.691 | **0.047** | not identifiable |

Supplementary Table 3: **Measurement-sufficiency evidence within individual hepatocyte production batches.** Each batch supplies its own held-out-compound folds, replicate pairs and reliability estimate. Values are medians over five folds, except reliability, which is estimated separately from replicate wells. Bold values fall outside the criteria for a quantitative proxy. Compound coverage differs among batches: batches 25, 26 and 27 contain roughly 630 compounds each, and batch 30 contains 286. Top-5% recall uses the top $k$ held-out compounds per fold, with $k = 7$ in the first three batches and $k = 3$ in batch 30. This limited resolution did not determine the classifications because magnitude-rank fidelity was below the proxy criterion in every batch.

| Readout | Batch | $r$ | $\rho$ | Var. | Recall | Reliab. | Tier |
|---|---|---|---|---|---|---|---|
| metabolic activity | 25 | 0.773 | **0.602** | 1.261 | **0.571** | 0.515 | unresolved |
| metabolic activity | 26 | 0.830 | **0.632** | 0.962 | 0.857 | 0.611 | unresolved |
| metabolic activity | 27 | 0.806 | **0.489** | 1.048 | 0.714 | 0.580 | unresolved |
| metabolic activity | 30 | 0.805 | **0.522** | 0.679 | 0.667 | 0.455 | unresolved |
| LDH release | 25 | **0.571** | **0.669** | 0.920 | **0.571** | 0.432 | measurement required |
| LDH release | 26 | **0.689** | **0.622** | 0.762 | 0.857 | 0.561 | unresolved |
| LDH release | 27 | **0.685** | **0.529** | 0.992 | **0.571** | 0.342 | unresolved |
| LDH release | 30 | **0.407** | **0.443** | 0.535 | **0.333** | 0.315 | measurement required |

Supplementary Table 4: **Sensitivity of anti-TOMM20 guide-partition reliability to gene-set composition.** Within-screen reliability pools gene–endpoint responses after standardising each endpoint by the observed spread of the full-guide response; the reported value is the median across screens. "Genes" reports the number retained after mapping to eligible screen genes. Each predefined set is compared with 200 size-matched random subsets from the same screen. "Outside control" indicates a value above the 95th centile of those draws. This comparison assesses gene-set composition at a matched set size; all reported reliability estimates remain below the operational criterion of 0.30.

| Gene set | Genes | Reliability | Size-matched random, median [90%] | Outside control |
|---|---|---|---|---|
| all genes | 20,213 | 0.047 | — | — |
| CEGv2 core essential[1] | 668 | 0.069 | 0.047 [0.038, 0.057] | yes |
| MitoCarta 3.0[2] | 1,066 | 0.051 | 0.048 [0.041, 0.054] | no |

Supplementary Table 5: **Sensitivity of reporter tiers to the decision thresholds on the common control-referenced scale.** Each of eight thresholds was varied independently in stricter and less stringent directions, first by 0.05 in the metric's units and then by 10% of the threshold value. The last row varies all eight simultaneously as a joint sensitivity analysis. Entries are the numbers of reporters whose tier changed among 52. The final column gives the range in the combined number of quantitative and ranking proxies; the primary analysis classified 37 reporters as proxies. Across all 36 scenarios, 28 reporters keep their exact tier and 40 keep their proxy or non-proxy status. Independent threshold changes altered a median of one reporter (maximum 8). The nine not-identifiable reporters were determined by target reliability: their number remained 9 whenever that threshold was unchanged and ranged from 6 to 14 when it was varied. Across all scenarios, the proxy count ranged from 30 to 42. Per-reporter assignments and transition matrices accompany the source data.

| | Moved by 0.05 | | Moved by 10% | | Proxy |
|---|---|---|---|---|---|
| Threshold moved | looser | stricter | looser | stricter | total |
| recoverability $\geq 0.70$ | 2 | 2 | 2 | 8 | 37 |
| magnitude Spearman $\geq 0.70$ | 2 | 3 | 2 | 6 | 31–39 |
| variance ratio lower edge 0.50 | 1 | 0 | 1 | 0 | 37 |
| variance ratio upper edge 1.50 | 0 | 0 | 0 | 0 | 37 |
| top-5% recall $\geq 0.60$ (quantitative) | 0 | 1 | 0 | 1 | 37 |
| top-5% recall $\geq 0.50$ (ranking) | 0 | 1 | 0 | 1 | 36–37 |
| reliability $\geq 0.30$ | 3 | 5 | 3 | 4 | 36–40 |
| recoverability $< 0.60$ (measurement required) | 3 | 0 | 3 | 0 | 37 |
| all eight together | 12 | 11 | 12 | 16 | 30–42 |

Supplementary Table 6: **Training and model selection for the ten full-label predictors.** All methods received the same training-standardized 172-dimensional phase representation, reporter-specific standardized target blocks and predefined train, validation and test assignments. "Observed only" means that unmeasured endpoints did not enter either the loss numerator or denominator. Specialist models were fitted independently for each reporter; shared models retained all source-reporter supervision available in a fold. Neural checkpoints were selected without access to outer-test targets. The table records validation-selected benchmark states; held-out-gene response replacement also uses the archived parameter-averaged neural checkpoints described in Methods. Architectures, optimization settings and software versions accompany the released analysis code.

| Method | Training scope | Implementation class | Target and loss semantics | Validation and model selection | Implementation and provenance |
|---|---|---|---|---|---|
| Ridge | Reporter-specific; one estimator per observed endpoint | Float64 closed-form ridge regression without an additional intercept | Each endpoint used its observed rows; validation and test inputs were clipped to the training feature range | $\alpha \in \{0.1, 1, 10, 100, 1000\}$ selected by validation MSE | Released study implementation; centred inputs and targets make the intercept redundant |
| GBDT | Reporter-specific; one estimator per observed endpoint | XGBoost regression trees[3]: 300 trees, depth 6, learning rate 0.05, row and column subsampling 0.8 | Squared-error objective on observed training rows | Validation RMSE with early-stopping patience 20 | Official XGBoost implementation; complete software environment and configuration released |
| CatBoost | Reporter-specific; one joint multi-output estimator per reporter | CatBoost[4]: 300 iterations, depth 6, learning rate 0.05, $L_2$ leaf penalty 3 and 128 borders | MultiRMSE over the complete observed reporter block; Plain boosting | Validation early stopping with patience 20 and restoration of the best iteration | Official CatBoost implementation; complete software environment and configuration released |
| MLP | Reporter-specific; one multi-output network per reporter | Hidden widths 256 and 128; Linear–GELU–LayerNorm blocks | Mean squared error over the standardized observed reporter block | AdamW, learning rate $10^{-3}$, weight decay $10^{-4}$, batch 1,024; up to 100 epochs, patience 10; lowest validation MSE restored | Released study implementation and machine-readable configuration |
| TabM | Reporter-specific; one multi-output model per reporter | Three 256-wide blocks, parameter-efficient ensemble size 16 and dropout 0.05[5] | Observed-only squared error over the reporter endpoint block; ensemble predictions averaged before export | AdamW, learning rate $10^{-3}$, weight decay $3 \times 10^{-4}$; lowest validation MSE restored | Official TabM implementation; source revision and configuration released |
| scButterfly | Reporter-specific paired phase–reporter model | Phase encoder 256–128, target encoder 128–128, latent dimension 128, dropout 0.1 and phase masking 0.5[6] | Paired reconstruction, translation, KL and adversarial objectives; unavailable endpoints masked | 100 phase-pretraining, 100 target-pretraining and 200 joint epochs; 50-epoch KL warm-up; best validation state restored | Public scButterfly implementation with released sparse-target adapter |
| ResMLP | Shared across source reporters with reporter-specific output heads | Width 512, expansion 1,024, four residual blocks, head width 128 and dropout 0.05 | Reporter-balanced observed-only MSE, equalized over endpoints and active reporters | AdamW, learning rate $3 \times 10^{-4}$, weight decay $10^{-4}$; 100 epochs, two-epoch warm-up, cosine decay and EMA 0.999 | Released study implementation and machine-readable configuration |
| MultiTab | Shared across the source-reporter panel with reporter-specific tokens and heads | Token dimension 32, four attention heads, two blocks, feed-forward width 128, head width 64 and dropout 0.05 | Reporter-balanced observed-only MSE as for ResMLP | AdamW, learning rate $3 \times 10^{-4}$, weight decay $10^{-4}$; 100 epochs, two-epoch warm-up, cosine decay and EMA 0.999 | Released study implementation and machine-readable configuration |
| MIDAS | Shared across all source reporters represented as sparse modalities | Biological latent 128, technical latent 32, modality and shared widths 768, two-layer decoder, dropout 0.1[7] | Phase and observed-only reporter reconstruction, biological and technical KL terms and modality alignment (weight 50) | AdamW, learning rate $10^{-4}$, weight decay 0.01; 100 joint epochs; lowest mean reporter validation MSE restored | Public MIDAS implementation with released sparse-modality adapter |
| scPair | Reporter-specific paired phase–reporter model | Encoder widths 256–64, cross-modal width 128 and dropout 0.1[8] | Sum of phase reconstruction, reporter reconstruction and bidirectional translation MSE on complete reporter rows | AdamW, learning rate $10^{-4}$, weight decay $10^{-4}$, batch 16,384; 40 epochs of 64 steps; best validation state restored | Public scPair modules with released paired-modality adapter; EMA and top-three checkpoint averages archived |

Supplementary Table 7: **Dependence of external measurement assignments on the numerical operating points.** Prediction and reliability metrics were held fixed while applying the same 32 single-threshold and four joint variations used in Supplementary Table 5. Entries count configurations retaining the baseline tier; the final column lists other tiers observed across the 36 variations. These counts describe the chosen configurations, not confidence probabilities. Q, quantitative proxy; R, ranking proxy; M, measurement required; N, not identifiable; U, unresolved. Main summaries and production-batch summaries are shown separately because their compound coverage and reference estimates differ. The low-reliability N assignment can remain unchanged while fidelity criteria change; all criterion-level comparisons accompany Supplementary Data 2.

| Measurement context | Baseline | Unchanged tier | | Other tiers |
|---|---|---|---|---|
| | | Single (/32) | Joint (/4) | |
| *Main applications* | | | | |
| OASIS metabolic activity | U | 32/32 | 4/4 | — |
| OASIS LDH release | U | 31/32 | 3/4 | Q/R |
| PERISCOPE anti-TOMM20 (cell) | N | 32/32 | 4/4 | — |
| PERISCOPE anti-TOMM20 (guide) | N | 32/32 | 4/4 | — |
| *OASIS batch applications* | | | | |
| OASIS metabolic activity, batch 25 | U | 32/32 | 4/4 | — |
| OASIS LDH release, batch 25 | M | 28/32 | 2/4 | R/U |
| OASIS metabolic activity, batch 26 | U | 31/32 | 3/4 | Q |
| OASIS LDH release, batch 26 | U | 32/32 | 4/4 | — |
| OASIS metabolic activity, batch 27 | U | 32/32 | 4/4 | — |
| OASIS LDH release, batch 27 | U | 31/32 | 3/4 | N |
| OASIS metabolic activity, batch 30 | U | 32/32 | 4/4 | — |
| OASIS LDH release, batch 30 | M | 30/32 | 2/4 | N |

Supplementary Table 8: **Calibration rules for task-use decisions.** All gates use calibration perturbations; evaluation perturbations supply the independent task outcome. Here $r$ is signed/profile recovery, $\rho$ is rank of response magnitudes, $v$ is predicted/measured magnitude variance, $h$ is top-budget recall, and $q$ is reference-repeat signed correlation. OPS full-response does not include $q$. The fixed response tiers in Fig. 3d are a separate atlas-wide diagnosis.

| Rule | Calibration gate | Application |
|---|---|---|
| Accept all | No gate | OPS, Cell Health, LINCS |
| Recovery | $r \geq 0.70$ | All three |
| Rank | $\rho \geq 0.70$ | All three |
| Rank + hit | $\rho \geq 0.70$ and $h \geq 0.50$ | OPS |
| Full response | $r, \rho \geq 0.70$, $0.50 \leq v \leq 1.50$, $h \geq 0.60$ | OPS; task-budget hit gate |
| Original-budget full | Same, with $h$ at 5% | OPS sensitivity |
| Task | $h \geq 0.60$ | Cell Health, LINCS |
| Task + repeat | $h \geq 0.60$, $q \geq 0.30$ | Cell Health, LINCS |
| Task + repeat + rank | $h \geq 0.60$, $q \geq 0.30$, $\rho \geq 0.70$ | LINCS |

Supplementary Table 9: **Decision units, partitions and random-selection references.** Candidate counts and budgets refer to the primary evaluation, not pooled observations across assays. Random expected recall is $k/N$ for selecting $k$ of $N$ candidates; it is not an acceptance threshold. Repeat estimates follow each assay's acquisition unit.

| | OPS | Cell Health | LINCS |
|---|---|---|---|
| Input / target | Phase features / fluorescent reporter profiles | Fluorescent morphology / cell-health readouts | Fluorescent morphology / landmark RNA |
| Paired unit | Same cell, aggregated by gene–screen | Guide within cell context; separate assay wells | Chemical condition; separate assay plates |
| Acquisition context | A549, matched phase/reporter images | ES2 and HCC44, separate health panels | A549, images at 48 h and RNA at 24 h |
| Decision case | Reporter–outer fold | Endpoint–cell context | RNA probe–predictor |
| Primary cases | 260 (52 × 5) | 20 (10 × 2) | 978 per predictor |
| Predictor | Fixed MLP predictions; three sensitivities | Fixed Ridge per case | Ridge and MLP |
| Training | Three of five original gene folds | 25 genes | 775 compounds |
| Model validation | Next cyclic fold | Fixed penalty | Training-only splits |
| Decision calibration | 91–111 genes per fold | 12 genes | 257 compounds |
| Evaluation candidates | 92–111 genes per fold | 12 genes | 260 compounds |
| Evaluation hits | $\lceil 0.20N \rceil$ | 4 | 26 |
| Required retained hits | $\lceil 0.60k \rceil$ | 3 | 16 |
| Random expected recall | $k/N \approx 0.20$ | 1/3 | 0.10 |
| Repeat evidence | Within-screen cell halves; sensitivity cohort | Two endpoint plates | Odd/even X-plate groups |
| Input requirement | Paired fluorescent targets for training | Stained images | Stained images |
| Allocation sensitivity | 20 allocations without refitting | 20 gene partitions | 10 Ridge chemical-group partitions |

#### Supplementary Data 1

**OPS response, decision and functional evidence.** Reporter–predictor criteria and tiers for all 52 reporters; calibration/evaluation gene assignments and complete decision ledgers; matched-coverage comparisons and reporter-cluster intervals; ensemble hit membership, ranked and significant enrichment lists, and fixed budget/background/tie sensitivities. Bundled verification code checks fixed-rule acceptance against the case ledgers, reconstructs primary decision counts from calibration/evaluation metrics, and recalculates enrichment probabilities and adjusted values from stored term-overlap counts.

#### Supplementary Data 2

**External measurement evidence and threshold sensitivity.** Full-precision metrics, baseline assignments and criterion-level changes across 36 threshold configurations for four external summaries and eight hepatocyte batch summaries, with reconstruction code.

#### Supplementary Data 3

**External task predictions and decision evidence.** OASIS held-out well predictions, signed compound losses, selected hits, source comparisons and uncertainty estimates; Cell Health guide/gene predictions, two-plate references, partitions and all 20 endpoint–context decisions over 20 fixed gene splits; LINCS primary frozen predictions, compound-group assignments, consensus and A/B RNA references, probe annotations, all 978-probe decision ledgers, ten Ridge partitions and joint compound-bootstrap results. The archive includes source identifiers, feature manifests and verification code.

#### Supplementary Methods

##### Predictor-specific measurement criteria and prediction-state sensitivity

We assessed predictor dependence from existing out-of-fold predictions for all ten methods, 52 reporters and five held-out-gene partitions. Reporter identity, endpoint order, gene–screen correspondence and observed response arrays were checked before comparison. The same common control scale in Eq. 4 was applied to every method; all method-specific target arrays agreed after alignment. For each method, recoverability was the equal-endpoint Pearson correlation within fold followed by the arithmetic mean over five folds. Response vectors were averaged equally across screens within each held-out gene, and their Euclidean magnitudes supplied rank fidelity, the predicted-to-observed variance ratio and recall of the strongest 50 of 1,000 genes. No observed or predicted top-50 boundary had tied magnitudes. Target reliability was reused unchanged from the measured responses.

The primary comparison used the same checkpoint states for the standalone predictors and the response-replacement ensemble, with data partitions, numerical thresholds and rule precedence held fixed. Among the 43 reporters with reliability at least 0.30, 20 were proxies under all ten methods, 6 were not proxies under any method and 17 changed across this boundary. Exact tiers were unchanged across methods for 9 of the 43; 15 other reporters changed only between quantitative and ranking proxies, and 2 only between measurement required and unresolved. The nine reporters below the reliability threshold were not included in these stability denominators.

Repeating the complete comparison with the single-model benchmark predictions gave 22 reporters supported as proxies by all ten methods, 6 supported by none and 15 crossing the proxy boundary. Ridge, GBDT, CatBoost, MLP, TabM and MultiTab each supported the same 37 proxy targets as the manuscript ensemble in this single-state comparison, although their quantitative and ranking assignments differed. With checkpoint-averaged predictions, Ridge, GBDT, CatBoost, MLP and MultiTab supported that same set; TabM supported 35. Both complete matrices are included in Supplementary Data 1.

We also separated two ensemble definitions. Measuring recoverability directly on the checkpoint-averaged median predictions gave 30 quantitative, 7 ranking, 3 measurement-required, 9 not-identifiable and 3 unresolved assignments. Replacing that recovery value with the median of the ten same-state method scores changed ATG101 from quantitative to ranking: ensemble recovery was 0.714, whereas the median method score was 0.697. Building the median ensemble from single-model predictions instead gave 31 quantitative and 6 ranking proxies, with the other counts unchanged. Only SRRM2 changed relative to the primary ensemble: its magnitude-variance ratio was 0.504 in the single-state ensemble and 0.495 in the checkpoint-averaged ensemble, on opposite sides of the fixed 0.50 boundary. These comparisons preserve the numerical cut-offs and expose changes close to a threshold without interpreting them as distinct biological classes. The predictor analysis recalculated the response-level criteria, including strong-hit recall, but did not repeat functional-term retention separately for every method.

##### Retrospective task-decision validation

The assessment object comprises target, predictor, representation, context, aggregation unit, task and tolerance. Response tiers describe fidelity on the atlas; calibration rules predict support for a specific task, and evaluation perturbations determine success. Every case records calibration metrics, an acceptance decision, evaluation hit recall and one of four outcomes. Support coverage, false-support fraction and success recall use the denominators defined in Methods; no accepted cases makes false-support fraction undefined. The complete rules and resource-specific budgets are given in Supplementary Tables 8 and 9. Missing metrics do not satisfy a gate, and missing evaluation evidence is reported outside outcome denominators.

**OPS.** Frozen outer-test predictions were divided by complete gene into decision-calibration and evaluation halves, without reusing model-validation genes. The fitted predictor, common NTC scale and five outer folds remained fixed. The primary MLP all-52 cohort comprised 260 reporter–fold cases; Ridge, scButterfly and the fixed ensemble supplied sensitivities. The full-response task rule combined calibration recovery, rank, variance and top-20% recall. It did not include reference reliability; a secondary cohort was restricted to the 43 reporters passing the atlas reliability gate. Twenty allocations changed calibration/evaluation gene membership without retraining. Supported-use gain counted extra accepted successes as a fraction of all cases; failure outcomes were reported separately.

Paired reporter-cluster bootstrap kept folds and all repeated allocations together. Calibration-only scores also selected exactly 25%, 50%, 75% or 100% of the reporter cohort within each fold, rounded up, with fixed identifier-based tie resolution (Supplementary Figure 6). At 75% matched coverage in the primary MLP analysis, rank minus recovery false support was $-0.0205$ (95% interval $-0.0585$ to 0.0114). At fixed thresholds, the corresponding difference was $-0.0085$ ($-0.0514$ to 0.0306), although supported-use gain was positive. These comparisons quantify successful-use coverage and conditional failure separately.

**Cell Health.** Ten endpoints in each of ES2 and HCC44 yielded twenty cases. Eligible guides had finite target observations on both reference plates; A549 and unresolved gene aliases were excluded according to mapping and reference coverage, before fitting. Gene assignments grouped guides across both cell contexts. The model used 25 training genes; twelve calibration and twelve evaluation genes defined four-hit tasks. Consensus targets averaged equally across the two independently acquired endpoint plates after plate-specific infected-control centring. Fluorescent morphology and health endpoints were paired by guide and context, not cell or well. The candidate set was twelve genes, the hit count four and random expected recall $4/12 = 1/3$.

Task and task-plus-repeat acceptance sets were identical in the primary split, not merely equal in size. Both selected five cases: three supported successes and two failures, with seven missed successes and eight true rejections. Of ten consensus-successful cases overall, six also passed against both reference plates. Across the twenty fixed allocations, adding repeat rejected five of 400 case–split instances, all evaluation failures. Comparing task-plus-repeat with recovery-only gave a lower failure fraction in eight allocations, the same fraction in four and a higher fraction in eight. These allocation counts characterize this dataset, not independent repetitions. Family-balanced summaries retain the five prespecified endpoint families and their correlated membership (Supplementary Data 3).

**LINCS.** Cell Painting at 48 h and L1000 at 24 h were matched by chemical identity and actual 10 $\mu$M condition, with at least two plates per assay. Each chemical group and every replicate belonged to one of training, calibration or evaluation. Predictor hyperparameters and training epochs used training-only model-validation folds. RNA reference A combined odd X-acquisition groups and B even groups; both consisted of independent plate acquisitions. The consensus included A and B, so the three comparisons share reference observations. Compound–plate means, rather than cross-assay well pairings, were averaged equally across plates.

The primary split used 775 training, 257 calibration and 260 evaluation compounds. All 978 landmark probes were retained, with ambiguous gene annotations identified explicitly. Top-10% selection yielded 26 evaluation hits; task success required at least sixteen. Task-plus-repeat used calibration A/B signed Pearson $\geq$ 0.30, while its repeat-supported evaluation outcome required success against each reference separately. Primary task and task-plus-repeat sets were identical for both predictors. Across ten fixed Ridge allocations the repeat gate rejected eight additional endpoints, seven failures and one success. Joint compound-group resampling preserved the entire probe panel and recomputed evaluation hit sets, with calibration acceptance fixed. Zero acceptance in rank-gated rules is reported with undefined false-support fraction.

The decision analyses were retrospective, with protocol settings fixed before execution and highlighted targets drawn from the complete outcome ledgers.

##### Significant and ranked functional evidence

Functional analyses used fixed ensemble predictions and the same response scale for all 52 reporters. The eligible enrichment background contained 1,000 knockout genes; NTC was excluded. Term size was 3–250 mapped assayed genes, and observed and predicted hit sets used the same annotation membership. Hypergeometric $P$ values were BH-adjusted over every eligible term within each reporter, family and hit budget, separately for observed and predicted lists. Significant sets used adjusted $P \leq 0.05$. Recall was undefined for empty measured sets and Jaccard undefined when both sets were empty. Ranked-list Jaccard instead compared fixed top-ten lists regardless of significance.

The cohort-relative downstream-retention index used 1,001 source-atlas labels, including NTC, and source-order resolution of ranking ties. The main functional panels instead use the 1,000-gene background and deterministic full-term-name ties. A sensitivity comparison found that changing only the background left all 156 primary ranked-list comparisons unchanged, whereas full-term-name tie resolution changed twenty. Complete enrichment scores and both sensitivity tables are included in Supplementary Data 1. Budget sensitivities used 5%, 10% and 20%; list-length sensitivities used 5, 10 and 20 terms. Redundancy-aware and member-matched summaries describe alternative ways of comparing term lists; they were not used for tier assignment.

At the primary budget, measured significant biological-process sets were nonempty for 33 reporters and empty for 19. Twenty-one quantitative proxies had nonempty measured sets, and all retained at least one significant term. The zero top-ten overlap for VPS35 occurred with zero significant terms in both measured and predicted lists. At the larger fixed budgets, shared significant counts were four of five and five of eight. This distinguishes ranked-list disagreement from retention of measured significant evidence.

**Sensitivity to the cross-fold response scale**

The prediction models standardized each reporter endpoint using the training cells of its outer partition. Endpoint-wise correlations are invariant to this scaling, but combining response vectors across partitions can change their relative endpoint weights because the five training-set standard deviations differ. The primary response-profile analyses therefore used the common control reference in Eq. 4: the pooled population standard deviation of unique non-targeting-control cells after subtraction of their physical-screen-specific control mean. Conversion used the saved training-fold target standard deviations and control cells, without model refitting. The five out-of-fold row sets were disjoint and exhaustive, and target transforms agreed across methods for every reporter and fold.

As a sensitivity analysis, we repeated the response-profile, counterfactual-replacement and tier calculations in the original fold-standardized coordinates. The primary common-scale analysis classified 30 reporters as quantitative proxies, 7 as ranking proxies, 3 as requiring measurement, 9 as not identifiable and 3 as unresolved. The fold-standardized sensitivity gave 30, 6, 3, 9 and 4, respectively. ER NCLN was the only reporter to change tier: it was unresolved on the fold-standardized scale and a ranking proxy on the primary common scale. All recoverability and reliability values were unchanged. The number of reporters with a predicted-to-observed magnitude-variance ratio below 0.5 remained 11, of which 10 retained magnitude-rank correlation above 0.5. Paired values and the code for the scale comparison accompany the source data.

**Sensitivity of the reporter tiers to the threshold values**

We assessed how strongly the atlas-derived tier assignments depended on the exact threshold values. Each of the eight thresholds was varied independently in both directions under two schemes: an absolute change of 0.05 in the metric's units and a relative change of 10% of the threshold value. All eight thresholds were also varied together, yielding 36 scenarios. Reporter evidence remained the same in every scenario. Because the variance ratio is evaluated within a two-sided interval, greater stringency moved both limits towards 1.0 and lower stringency moved them apart. The original thresholds on the primary common control scale reproduced the five reported tier counts (30, 7, 3, 9 and 3).

The ranking-proxy rule is evaluated before the measurement-required rule. Thus, a reporter that preserves magnitude ranking and strong-hit recall is not assigned to measurement required solely because its recovery is below 0.60; the same precedence is retained throughout the threshold sensitivity analysis.

**Threshold sensitivity in the independent applications**

We applied the same 36 configurations to the four main external evidence summaries and eight OASIS production-batch summaries (Supplementary Table 7). Fidelity metrics retained the five-fold median aggregation used in the main analysis, and reliability retained the resource-specific reference estimate. We recalculated assignments from full-precision source values rather than the rounded table entries, without refitting predictors or changing the metric definitions. Alongside tier changes, we recorded changes in the four quantitative-fidelity conditions—recovery, magnitude rank, the two-sided variance interval and strong-hit recall—independently of the reliability gate. The analysis uses the same threshold variations as in OPS to assess sensitivity to the operating points, rather than calibrating thresholds for a new application.

The pooled OASIS metabolic-activity assignment remained unresolved in all 36 configurations. LDH release changed from unresolved to ranking proxy when only the rank threshold was reduced to 0.63, and to quantitative proxy under the joint less stringent relative variation; the other 34 configurations retained its baseline tier. Both PERISCOPE assignments remained not identifiable, although at least one quantitative-fidelity criterion changed whether it was met in 7 of 36 cell-derived and 4 of 36 guide-profile configurations. Thus, individual fidelity conditions could change while the reliability-gated classification stayed the same. Batch-specific changes are retained in Supplementary Table 7.

**Endpoint aggregation in the independent applications**

For the CellProfiler-based OASIS analysis, each biochemical readout contributed one primary endpoint. Recoverability was Pearson correlation between observed and predicted compound–plate means of the deposited plate-normalized endpoint within an outer fold, with control compounds excluded. For magnitude-rank fidelity, variance ratio and hit recall, these means were first centred on the corresponding within-fold plate-control mean and then averaged equally across plates for each held-out compound. Each reported fidelity value was the median over the five folds.

For PERISCOPE, recoverability was Pearson correlation pooled across held-out gene–plate–endpoint response coordinates within each outer fold. Before pooling, each endpoint's observed and predicted responses were divided by the standard deviation of its observed response coordinates in that fold. Endpoints were retained only when this standard deviation was positive and exceeded $10^{-3}$ times the median endpoint standard deviation. This pooled, endpoint-standardized statistic differs from the equal-endpoint mean correlation used for OPS in Eq. 2. For response-magnitude analyses, gene responses were first averaged equally across plates and endpoints were standardized by their observed across-gene standard deviations within fold before calculating the Euclidean norm. These evaluation scales were applied identically to observations and predictions and were not used for model fitting. The numerical decision thresholds were retained from OPS, while aggregation and replicate definitions reflected the available measurements in each resource.

**Per-batch analysis of the hepatocyte screen**

We repeated the OASIS analysis separately within each of the four production batches to assess variation in the evidence supporting substitution. Held-out-compound folds, replicate halves and reliability estimates were constructed within batch. Every retained compound had at least two wells within its batch, allowing split-half reliability to be estimated in all four batches. Reliability ranged from 0.31 to 0.61, compared with pooled estimates of 0.72 and 0.64 for the two reporters. The batch-specific and pooled analyses differ in replicate coverage and compound composition, so this comparison does not isolate the source of the reliability difference.

Top-5% recall used the top $k$ held-out compounds in each fold: $k = 11$ in the pooled analysis, $k = 7$ in batches 25–27 and $k = 3$ in batch 30. In the last batch, recall can therefore take only the values 0, 1/3, 2/3 and 1. This coarse resolution limits comparisons of hit retention, but did not determine the tiers because magnitude-rank fidelity remained below 0.70 in all batch–reporter combinations. Metabolic activity was unresolved in every batch. LDH release was unresolved in batches 26 and 27 and required measurement in batches 25 and 30, where recoverability was below 0.60 (Supplementary Table 3).

**Metabolic-activity prioritization: repeated measurements, controls and uncertainty**

The target was the normalized RealTime-Glo MT signal supplied in the OASIS metadata column `Metadata_mtt_normalized`. Rankings used the negative of the signed, plate-control-relative response, not its absolute magnitude. Within-compound aggregation matched actual concentrations before averaging sources and then the eight concentration positions. All 217 held-out compounds had eight positions across the union of available sources. Ninety-four had all eight matched concentrations in both sources: 38 were paired between sources 26 and 27, and 56 between sources 26 and 30. The other 123 comprised 31 source-25/26 and 92 source-25/27 pairs, with incomplete source-25 dose coverage. Complete-source results therefore describe the 94-compound subset rather than repeatability of the entire test population. Both cross-source directions and the four actual source-pair directions are reported in Supplementary Data 3; source roles were assigned by lexical order within compound.

The primary Ridge model and a fixed histogram gradient-boosting model each used all 768 features from the released brightfield DINO representation. Gradient boosting used 100 iterations, learning rate 0.1, at most 15 leaves, L2 regularization 10 and no early stopping. Diagnostic comparators used either same-plate-control-centred cell counts or acquisition metadata, with the same two model specifications and training partition. Counts came from fluorescence-derived segmentation. Metadata comprised numeric row, column and dose position, plus one-hot-encoded plate and production source, with training-only category fitting and unknown-category handling; compound identities and target values were excluded. DMSO dose position was set to zero. At the primary budget in the 217-compound population, brightfield Ridge and gradient boosting each recovered 10/11, cell-count Ridge and gradient boosting recovered 8/11 and 7/11, and metadata models recovered 0/11 and 1/11. These comparisons describe available abundance and acquisition information; they do not isolate target-specific mechanisms or a conditional contribution beyond cell count. For the primary brightfield model, the selected set's observed mean loss was 0.3336 (range, 0.2151–0.8995), compared with 0.3377 for the measured top eleven and a median of $-0.0008$ across all 217 compounds. Units are changes in normalized signal, not percentages of viable or dead cells.

For each budget, observed and predicted top sets had the same size, $k = \lceil fN \rceil$, so their overlap divided by $k$ was both recall and precision. Compound identifiers resolved ties deterministically. Bootstrap resampling drew compounds 2,000 times, retaining each compound's doses and source roles, assigning unique instance identifiers to repeated draws and recomputing the top sets at each draw. Paired differences used the same draws for prediction-to-other-source and measured-source-to-other-source recall. The primary Ridge recall interval was 0.545–1.000 for all 217 compounds and 0.400–1.000 for each of the three pooled or directional 94-compound comparisons. Cross-source differences relative to measured repeats were zero at the point estimate, with intervals $-0.60$ to 0.40 and $-0.40$ to 0.20 for the two directions. These intervals condition on fitted models and observed production sources; they do not estimate uncertainty for new donors or new sources. No equivalence or non-inferiority margin was specified.

We also disrupted test prediction–compound correspondence within plate for each fixed brightfield model, using 2,000 permutations. Dose position was constant within each plate, so plate–dose strata reduced to 61 within-plate groups; all test wells were movable. This preserved the acquisition structure while breaking compound correspondence without refitting a model. For Ridge at the primary budget, mean null recall was 0.0538 and the central 95% of the permutation distribution was 0–0.1818, compared with observed recall of 0.9091. These controls apply to the full 217-compound analysis only. Exact hypergeometric random-selection references were calculated separately for each population and budget; their intervals describe random overlap, not model-performance uncertainty. Fixed 10% and 20% budgets recovered 17/22 and 34/44 in the full population. All predictors, budgets and source comparisons are retained in Supplementary Data 3.

**Sensitivity of guide-partition reliability to endpoint scaling and gene selection**

The low guide-partition reliability of anti-TOMM20 was decisive for both PERISCOPE classifications. This statistic measures agreement between guide-defined perturbation responses, combining measurement, guide and biological variation. We therefore examined its sensitivity to endpoint scaling and gene-set composition. Within each plate, reliability was calculated by pooling the paired gene–endpoint responses from disjoint sgRNA halves after dividing each endpoint by the observed spread of the full-guide response. Plate-specific correlations were then summarized by their median. As a scaling comparison, pooling in native units gave 0.050 across all genes and a median of 0.102 across 200 random 668-gene subsets; the endpoint-standardized estimates were 0.047 for both the complete set and the median subset. Standardization reduced the sensitivity of the pooled statistic to this change in gene sampling.

We next evaluated two externally defined gene sets selected independently of target values. The source sets contained 684 core essential genes from CEGv2[1] and 1,136 mitochondrial genes from MitoCarta 3.0[2]; after intersection with eligible screen genes, 668 and 1,066, respectively, entered the analysis. Each set was compared with 200 size-matched random subsets from the same screen (Supplementary Table 4). Reliability was 0.069 for core essential genes and 0.051 for mitochondrial genes. Only the essential-gene estimate exceeded the 95th centile of its size-matched reference distribution, and both remained below 0.30. Thus, the low reliability persisted under the two tested gene selections, although gene-set composition affected its magnitude.

**Leave-one-dye-out substitution**

To examine prediction between fluorescent channels, we predicted each of the four Cell Painting dyes from the other three at the guide-profile level, using the same holdout design and decision thresholds. Features involving the held-out dye were excluded from the input, including cross-channel features shared with a predictor dye. All analyses used the deposited masks, whose segmentation included the target channel.

**Permuted-input negative control**

For the CellProfiler-based OASIS and PERISCOPE controls, we permuted input profiles among observations within the same plate, retaining target measurements, perturbation labels and control status. This preserves plate membership and the marginal input distribution while disrupting the input–target correspondence used for prediction. OASIS profiles were permuted at the well level and PERISCOPE profiles at the guide-profile level. The OASIS model retained its fixed Ridge penalty of $\alpha = 1$. For PERISCOPE, the permuted-input analysis used a fixed penalty-to-training-row ratio of $\alpha/n = 0.01$, the value selected in four of the five main-analysis folds; the penalty was not reselected after permutation. Supplementary Table 2 reports the nested-selected main analysis, whereas Supplementary Note 1 reports the fixed-penalty permutation. Reliability was calculated from the unchanged target measurements and was therefore identical before and after input permutation.

#### Supplementary Note 1: Input permutation distinguishes prediction failure from inadequate target reproducibility

Within-plate permutation of OASIS CellProfiler-derived brightfield profiles reduced recoverability from 0.83 to 0.09 for metabolic activity and from 0.67 to 0.05 for LDH release. Magnitude-rank agreement approached zero, the predicted-to-observed magnitude-variance ratio was approximately 0.02, and strong-hit recall was low. Target reliability was unchanged. Both reporters were consequently classified as measurement required: their measured responses were reproducible, but the permuted morphology profiles did not recover them.

For PERISCOPE, the guide-profile permutation gave recoverability of $-0.0004$, magnitude-rank correlation of 0.022, variance ratio of 0.025 and strong-hit recall of 0.064 under the fixed-penalty control described above. The anti-TOMM20 classification remained not identifiable because guide-partition reliability was unchanged at 0.047, below the initial reliability criterion. Together, the two controls separate loss of input–target information from insufficient reproducibility of the reference response (Supplementary Fig. 1b).

#### Supplementary Note 2: Low guide-partition reliability persists across the Cell Painting dye targets

Predicting each Cell Painting dye from the other three gave guide-partition reliabilities of 0.032–0.054, all below 0.30 (Supplementary Fig. 1b). All four were therefore classified as not identifiable, as was anti-TOMM20, although recoverability ranged from 0.45 for F-actin to 0.80 for the endoplasmic reticulum. This shared classification reflects limited reference reproducibility, not uniform predictive performance. The reliability limitation thus extends beyond the antibody endpoint in this dataset. These comparisons do not distinguish guide efficacy, cell sampling, measurement variation or biological heterogeneity as its source.
